# A texture statistics encoding model reveals hierarchical feature selectivity across human visual cortex

**DOI:** 10.1101/2022.09.23.509292

**Authors:** Margaret M. Henderson, Michael J. Tarr, Leila Wehbe

## Abstract

Mid-level visual features, such as contour and texture, provide a computational link between low- and high-level visual representations. While the detailed nature of mid-level representations in the brain is not yet fully understood, past work has suggested that a texture statistics model (P-S model; Portilla and Simoncelli, 2000) is a candidate for predicting neural responses in areas V1-V4 as well as human behavioral data. However, it is not currently known how well this model accounts for the responses of higher visual cortex regions to natural scene images. To examine this, we constructed single voxel encoding models based on P-S statistics and fit the models to fMRI data from human subjects (male and female) from the Natural Scenes Dataset (Allen et al., 2021). We demonstrate that the texture statistics encoding model can predict the held-out responses of individual voxels in early retinotopic areas as well as higher-level category-selective areas. The ability of the model to reliably predict signal in higher visual cortex voxels suggests that the representation of texture statistics features is widespread throughout the brain, potentially playing a role in higher-order processes like object recognition. Furthermore, we use variance partitioning analyses to identify which features are most uniquely predictive of brain responses, and show that the contributions of higher-order texture features increases from early areas to higher areas on the ventral and lateral surface of the brain. These results provide a key step forward in characterizing how mid-level feature representations emerge hierarchically across the visual system.

**Significance Statement:** Intermediate visual features, like texture, play an important role in cortical computations and may contribute to tasks like object and scene recognition. Here, we used a texture model proposed in past work to construct encoding models that predict the responses of neural populations in human visual cortex (measured with fMRI) to natural scene stimuli. We show that responses of neural populations at multiple levels of the visual system can be predicted by this model, and that the model is able to reveal an increase in the complexity of feature representations from early retinotopic cortex to higher areas of ventral and lateral visual cortex. These results support the idea that texture-like representations may play a broad underlying role in visual processing.

## Introduction

Information in visual cortex is processed by a series of hierarchically organized brain regions, with the complexity of representations increasing at each level. While there are intuitive explanations for response properties at the ends of this hierarchy (e.g., oriented spatial frequency filters in primary visual cortex (Carandini et al., 2005; Hubel & Wiesel, 1962) or object and category representations in inferotemporal cortex (Desimone et al., 1984; Grill-Spector & Weiner, 2014)), the computations performed at intermediate levels have proven more challenging to describe. These intermediate- or mid-level visual areas are thought to represent features like contour and texture, which play an important role in figure-ground segmentation, shape processing, and object and scene classification (Bergen & Landy, 1991; Connor et al., 2007; Peirce, 2015; Ullman et al., 2002; Walther & Shen, 2014). Thus, developing a robust model of mid-level representation is fundamental for understanding how the visual system extracts meaningful information from the environment.

Computational texture models, particularly the influential model proposed by Portilla and Simoncelli (Portilla and Simoncelli, 2000; hereafter referred to as the “P-S model”), have proven useful in understanding the mid-level features that drive visual responses. The P-S model is constructed using a steerable pyramid decomposition (Simoncelli & Freeman, 1995) to extract features at different spatial scales and orientations, and includes features at multiple levels of complexity, including simple luminance and spectral statistics, as well as higher-order correlation statistics (Figure 1). The P-S model can account for aspects of texture sensitivity in primate visual areas such as V1, V2, and V4 (Freeman et al., 2013; Hatanaka et al., 2022; Okazawa et al., 2017, 2015), with sensitivity to the higher-order model features increasing from V1 to V2 to V4 (Freeman et al., 2013; Hatanaka et al., 2022; Okazawa et al., 2017). In humans, functional magnetic resonance imaging (fMRI) studies suggest that P-S statistics can account for some response properties in early visual and ventral visual cortex (Baumgartner & Gegenfurtner, 2016; Long et al., 2016). The P-S model also captures key aspects of human behavior, including perceptual discrimination of texture patches (Balas, 2006; Freeman & Simoncelli, 2011), crowding (Balas et al., 2009), search performance (Rosenholtz et al., 2012) and discrimination of high-level object properties such as real-world size and animacy (Long et al., 2016; Long et al., 2017).

**Figure 1:**
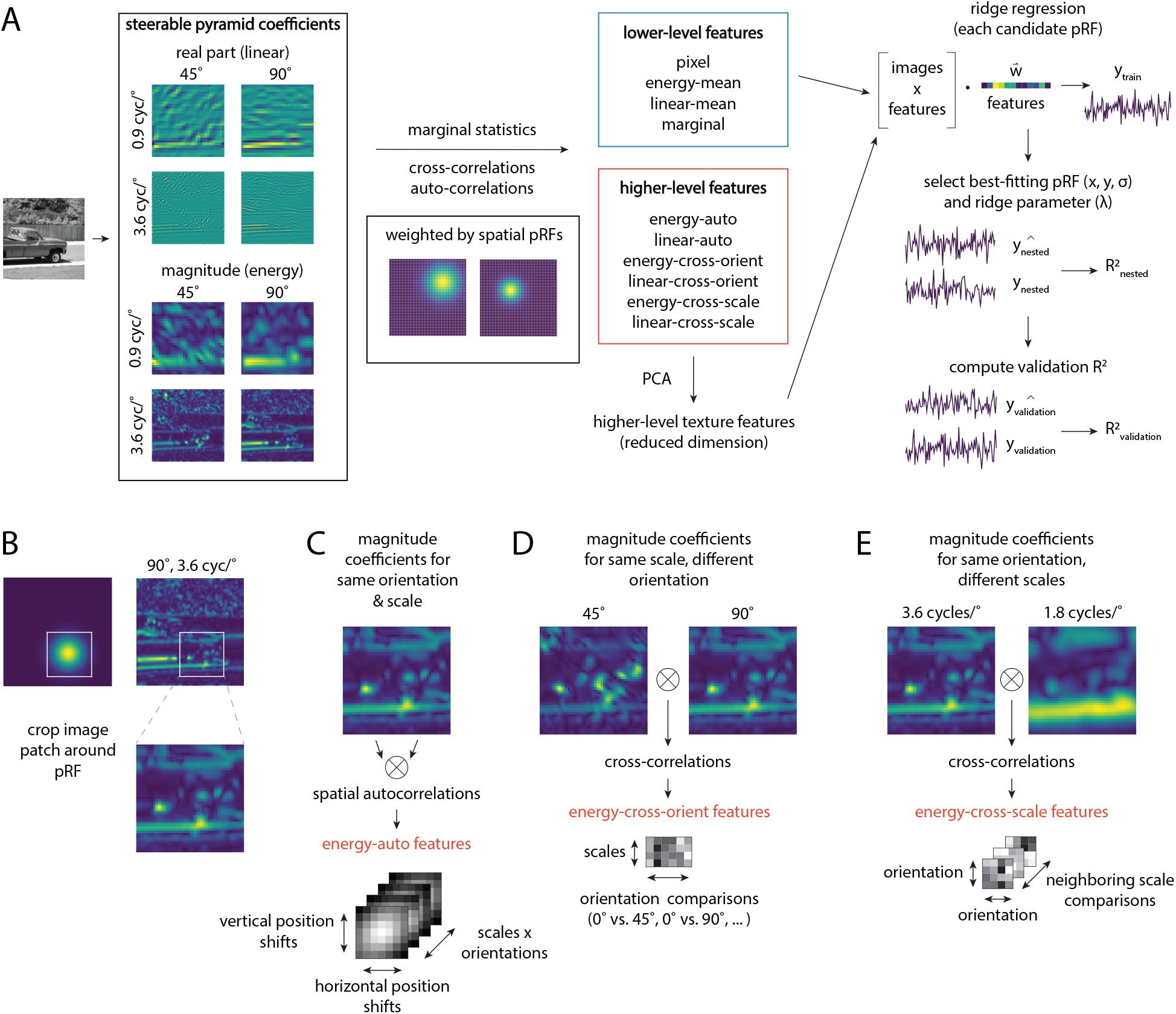
**(A)** Overview of the texture statistics model (Portilla & Simoncelli, 2000) and voxelwise encoding model fitting procedure. For each image, we first use a steerable pyramid (Simoncelli & Freeman, 1995) to decompose the image into four orientation and four frequency bands (note that only 2 orientations and frequencies are shown here for illustration). We then use the steerable pyramid representation to extract lower-level texture features, which include marginal statistics of the pyramid coefficients, and higher-level texture features, which consist of higher-order correlations of the pyramid coefficients (see *Methods: Texture statistics features* for details). Each of these features is computed for each population receptive field (pRF) in a grid of candidate pRFs, using the pRF as a spatial weighting matrix. Before fitting the model, we reduce the dimensionality of the higher-level texture features using PCA. We then use ridge regression to fit a set of weights for each voxel that predict its response to each image as a weighted sum of all the texture statistics features extracted from that image. We perform this fitting separately for each candidate pRF, and use the loss on a held-out (nested) data subset to choose the best pRF. Finally, we compute the model’s accuracy (*R*^2^) on a held-out validation set. See *Methods: Model fitting procedure* for more details. **(B-E)** Illustrations of how the higher-level model features are computed based on coefficients of the steerable pyramid, for one example pRF. The image is cropped to yield an image patch spanning *±* 2 *σ* from the pRF center; the pRF is cropped in the same way and used as a weighting matrix. **(C)** The *energy-auto* features are computed by correlating each magnitude feature map with spatially-shifted versions of itself (i.e., autocorrelations). **(D)** The *energy-cross-orient* features are computed by cross-correlating magnitude feature maps corresponding to the same scale but different orientation. **(E)** The *energy-cross-scale* features are computed by cross-correlating magnitude feature maps corresponding to different scales, after up-sampling the map corresponding to the coarser scale. The *linear-auto, linear-cross-orient*, and *linear-cross-scale* features are computed similarly, but using the real parts of steerable pyramid coefficients. See Table 1-1 for the total number of features included in each subset.

Despite these promising results, it is not yet known how well the P-S model captures neural responses to natural scene images, particularly in higher visual cortex. The majority of past studies have used synthetic stimuli or textures that are relatively homogeneous across space (e.g., Freeman and Simoncelli, 2011; Okazawa et al., 2017, 2015). In contrast, natural scenes, which include objects and other complex, spatially localized elements, may prove more challenging to characterize (Portilla & Simoncelli, 2000). One recent study did examine the P-S model’s prediction of neural responses in macaque V1 and V4 for naturalistic stimuli (Hatanaka et al., 2022); however, this study did not consider higher visual cortex. Under the hypothesis that mid-level features also play a key role in the neural processing of object and scene categories (Bracci et al., 2017; Connor et al., 2007; Groen et al., 2017; Long et al., 2018; Nasr et al., 2014), modeling responses to natural images within higher visual cortex is a critical test of the P-S model’s ability to generalize to real-world vision.

To address this question, we developed a forward encoding model (Naselaris & Kay, 2015; Serences & Saproo, 2012) that uses P-S model texture statistics to predict voxelwise responses from the Natural Scenes Dataset (NSD; Allen et al., 2021), and used variance partitioning to probe the contributions made by each subset of texture model features. Critically, our model: (1) achieves accurate predictions of held-out voxel responses throughout the visual hierarchy, with highest performance in early and mid-level areas; (2) recovers the hierarchical organization of visual cortex, with retinotopic areas best described by lower-level features and anterior category-selective areas best described by higher-level features. These and other results we present here facilitate a better understanding of how hierarchical computations give rise to mid-level representations in the brain, and how mid-level feature selectivity may be used to predict the emergence of a higher-level semantic representational space.

## Methods

### Acquisition and preprocessing of fMRI data

We utilized the Natural Scenes Dataset (NSD), a large-scale, publicly available fMRI dataset. Full details on the acquisition of this data can be found in Allen et al., 2021. The NSD includes whole-brain BOLD fMRI measurements from 8 human subjects (both male and female) who each viewed between 9,000-10,000 natural scene images over the course of a year (between 30-40 scan sessions). Functional scanning was conducted at 7T using whole-brain gradient-echo EPI at 1.8 mm resolution and 1.6 second repetition time (TR). Images were sampled from the Microsoft Common Objects in Context (COCO) database (Lin et al., 2014). Over the course of the experiment, each image was viewed approximately 3x, for a total of roughly 30,000 trials per subject (fewer for some subjects who did not complete the entire experiment). Of the approximately 10,000 images viewed by each subject, *∼*9,000 images were seen only by that subject, and 907 were viewed at least once by each subject. Each image was presented in color, at a size of 8.4° x 8.4° (° = degrees of visual angle), and was viewed for a duration of 3 seconds, with 1 second between trials. Throughout each scan, subjects performed a task in which they reported whether each image was new or old (i.e., whether or not it had been presented before in any session), while fixating centrally on a small fixation dot superimposed on each image.

As described in Allen et al., 2021, the functional data from all sessions were pre-processed using temporal interpolation to correct for slice timing differences and spatial interpolation to correct for head motion (see Allen et al., 2021 for details). All analyses were performed in each subject’s native volumetric space (1.8 mm resolution voxels). A general linear model was used to estimate beta weights for each voxel and each individual trial (Prince et al., 2022). We obtained the beta weights from Allen et al., 2021 after this stage, and we then performed a few additional steps to prepare the beta weights for our analyses. First, beta weights for each voxel were z-scored across all trials of each scan session. To improve the signal-to-noise ratio of the data, we then averaged the beta weights for each voxel across trials where the same image was shown (approximately three trials/image), resulting in a single value for each voxel in response to each of the unique (*∼*10,000) images. Note that for subjects who did not complete the entire experiment, there were fewer than 10,000 images.

We masked out a broad region of interest in visual cortex to include in all analyses. This region included voxels that were part of the “nsdgeneral” ROI described in Allen et al., 2021, which is meant to capture the general spatial extent of voxels that were responsive to the NSD image stimuli. To broaden the scope of brain areas included in our analyses, we additionally included voxels belonging to any ROI within several sets of ROI definitions: any ROI in the probabilistic atlas provided in Wang et al., 2015, which includes regions of the intraparietal sulcus, any ROI belonging to an early retinotopic area based on pRF mapping, and any face-selective, body-selective, or place-selective ROI identified through a functional category localizer task (see *Methods: Defining regions of interest (ROIs)* for details on pRF and category localizer tasks). We further thresholded voxels for inclusion according to their noise ceiling, which is a measure that captures the proportion of voxels’ response variance that can theoretically be explained by properties of the stimulus (Allen et al., 2021; Wu et al., 2006), using a threshold of 0.01. This large mask defines the extent of voxels that are included in whole-brain surface maps (e.g., Figure 3A). We also computed summary statistics at the ROI level using more fine-grained definitions of individual ROIs (e.g., Figure 3B), see *Methods: Defining regions of interest (ROIs)* for details.

**Figure 2:**
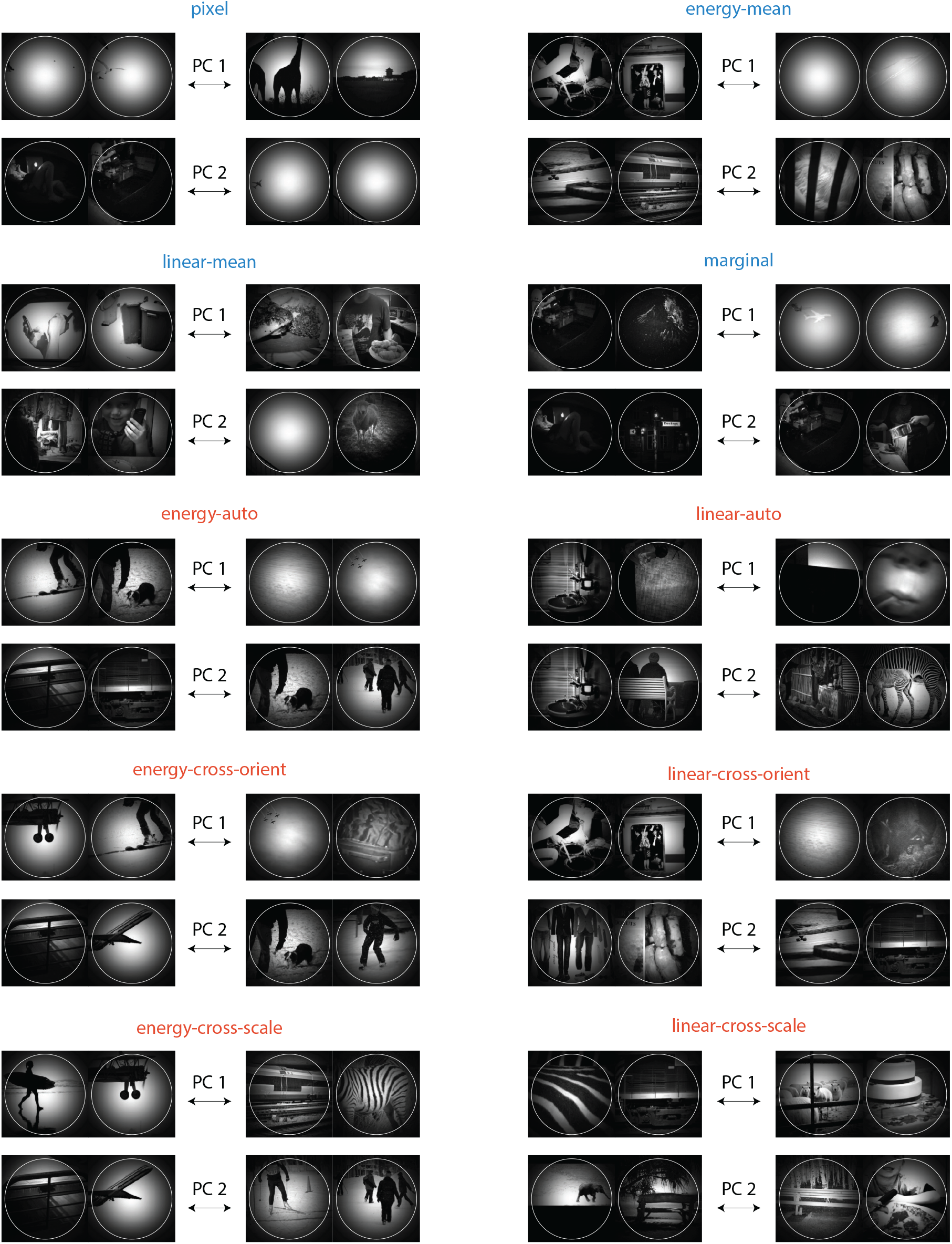
Visualization of the kinds of natural image features that are captured by each subset of P-S statistics, shown for one example pRF (pRF size = 1.48°). Each sub-panel of 8 images corresponds to one feature subset. To generate this visualization, we performed PCA on the model features corresponding to each feature subset, and identified the two most activating (left) and least activating (right) images for each of the first two principal components. To aid visualization of the image area that most strongly contributed to computing the features, we weighted each image according to the Gaussian profile of the pRF, and cropped the image to a square centered on the pRF. The white circle indicates a radius of *±* 2 *σ* from the pRF center. These visualizations are generated using only the images shown to S1, but similar results are obtained when using other sets of images.

**Figure 3:**
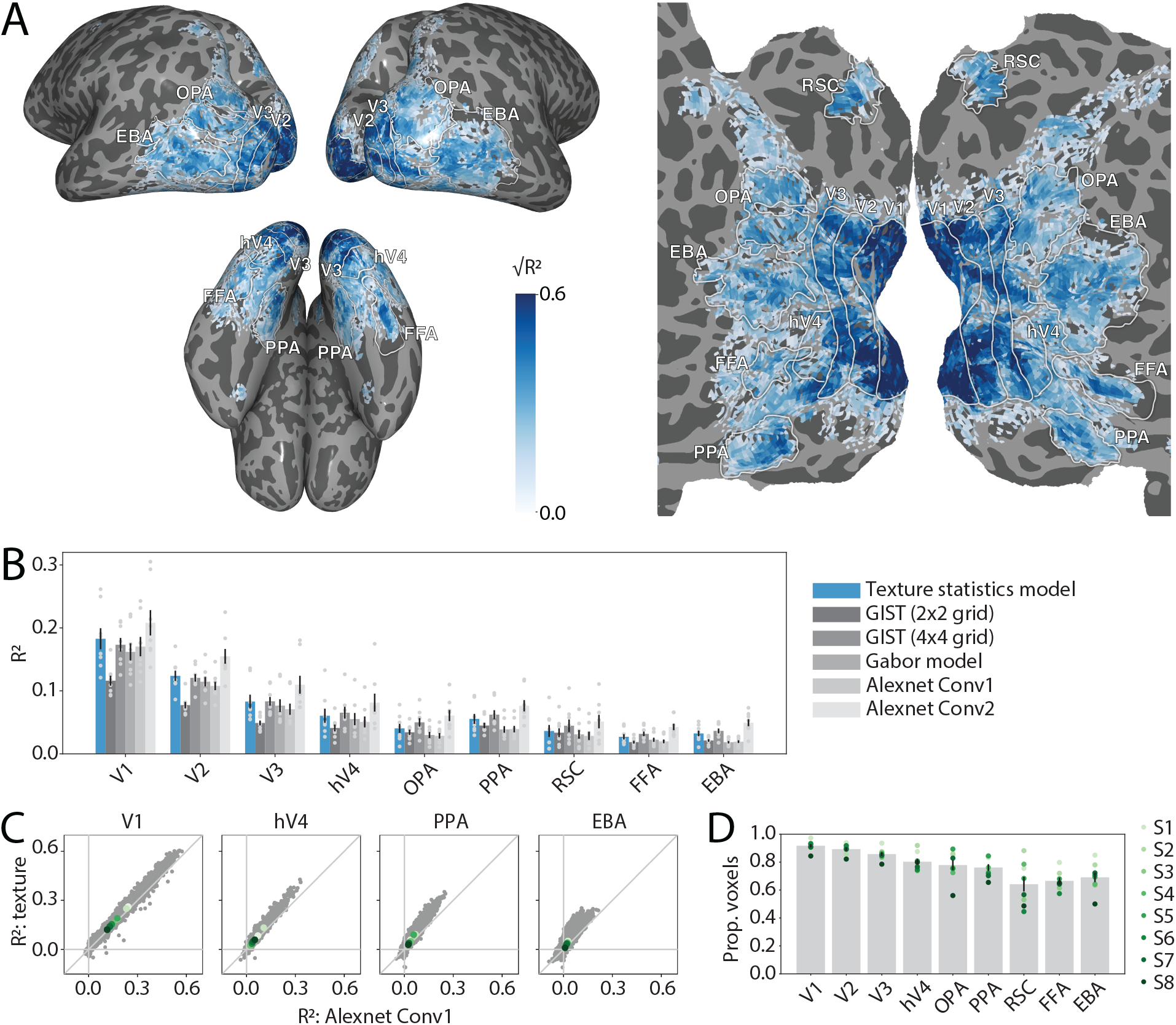
The texture statistics encoding model accurately predicts voxel responses across visual cortex, with highest accuracy in early and mid-level areas. **(A)** Model performance is shown on an inflated cortical surface (left) or flattened surface (right) for one example NSD subject, S1 (for the full set of 8 subjects, see Figure 3-1). *R*^2^ was computed using responses to an independent set of images that were not used to fit the model, see *Methods: Model fitting procedure* for details. For visualization purposes, values of *R*^2^ have been transformed to 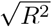, as this makes it easier to visually observe differences between colormap values. White outlines and labels indicate the location of functionally-defined retinotopic and category-selective ROIs in each subject; see *Methods: Defining regions of interest (ROIs)* for details. **(B)** Summary of *R*^2^ for our texture statistics model alongside several alternative low-level visual models (see *Methods: Alternative low-level visual models* for details). Bar heights and errorbars reflect mean *±* 1 SEM across subjects, light gray dots represent individual subjects. **(C)** *R*^2^ for the texture statistics model is plotted versus *R*^2^ for Alexnet Conv1, for four selected ROIs. Each gray dot represents a single voxel, dots in shades of green indicate the mean *R*^2^ for each individual subject (legend on far right). **(D)** Proportion of voxels in each ROI where validation set *R*^2^ for the texture model was significantly higher than chance, evaluated using a permutation test (see *Methods: Permutation testing* for details). Dots in shades of green indicate single subjects, bar heights and errorbar represent the mean *±* 1 SEM across subjects.

### Defining regions of interest (ROIs)

We defined retinotopic and category-selective ROIs based on functional localizers that were collected as part of the NSD experiment. A category localizer task (Stigliani et al., 2015) was used to define place-selective regions (parahippocampal place area/PPA, occipital place area/OPA, and retrosplenial cortex/RSC), a face-selective region (fusiform face area/FFA; we combined FFA-1 and FFA-2 into a single FFA region), and a body-selective region (extrastriate body area/EBA). A pRF mapping task (sweeping bar stimuli; Benson et al., 2018) was used to define early retinotopic visual ROIs V1, V2, V3, and hV4 (see Allen et al., 2021 for more details). When retinotopic and category-selective ROIs were overlapping with one another, we excluded any voxels that were overlapping from the retinotopic definition and added them to the corresponding category-selective ROI only. To ensure that the final set of category-selective ROIs were non-overlapping, we always prioritized face-selective ROIs over place- and body-selective ROIs, and prioritized place-selective ROIs over body selective ROIs.

### Texture statistics encoding model

#### Overview

Our encoding model incorporated parameters for each voxel’s feature selectivity as well as its spatial selectivity (St-Yves & Naselaris, 2017), see Figure 1 for an illustration. The inclusion of an explicit population receptive field (pRF) for each voxel is an aspect in which our model improves upon past work (Hatanaka et al., 2022; Okazawa et al., 2015), allowing us to adaptively fit voxel responses with a wide range of receptive field positions and sizes. During model fitting, both the pRF parameters and the texture feature weights are optimized simultaneously (see *Methods: Model fitting procedure* for details). To fit the model, we first created a grid of candidate population receptive fields or pRFs (see *Methods: Population receptive fields (pRFs)*), and extracted texture statistics features (see *Methods: Texture statistics features*) within each candidate pRF. For each pRF, we then used regularized regression to fit a linear model that predicts the voxel’s activation as a function of the texture statistics features corresponding to that pRF. The final encoding model for each voxel was computed by identifying the best-fitting model over all candidate pRFs. More detail on each step is provided in the following sections.

#### Population receptive fields (pRFs)

We modeled each candidate population receptive field (pRF) as a 2-dimensional Gaussian (Dumoulin & Wandell, 2008; St-Yves & Naselaris, 2017). Each pRF has three parameters, *x*_*0*_, *y*_*0*_, and *σ*, where [*x*_*0*_, *y*_*0*_] and *σ*, respectively, indicate the center and standard deviation of the 2-dimensional Gaussian response profile:

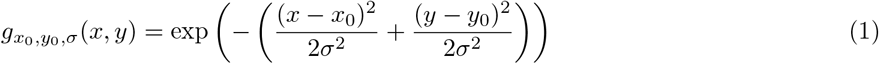

Our grid of candidate pRF parameters was designed to approximate a log-polar grid, where the spacing between adjacent pRF centers is linear in terms of polar angle position (*θ*), and nonlinear in terms of eccentricity (*r*), such that candidate centers were more closely spaced closer to the center of the visual field. This nonlinear eccentricity spacing was intended to account for the cortical magnification factor in human visual cortex, where the neuronal sampling of visual space is more dense close to the fovea (Duncan & Boynton, 2003). Specifically, we used 16 candidate polar angle positions that were linearly spaced, ranging from 0° to 337.5° in steps of 22.5°, and 10 candidate eccentricities that were logarithmically spaced, ranging from 0° to 7°. For each of these candidate centers, we generated 10 candidate *σ* values that were also spaced logarithmically, ranging from 0.17° to 8.4°. The complete grid of pRF parameters over all combinations of *r, θ*, and *σ* resulted in 1,600 pRFs.

Since our eccentricity range extended slightly beyond the physical image extent (an 8.4° square), some of the smaller pRFs at the largest *r* values were entirely non-overlapping with the image region (in contrast, larger pRFs at these large eccentricities were still partially overlapping with the image region). To address this issue, we removed from the grid any pRFs whose rough spatial extent (center *±σ*) was non-overlapping with the image region. This resulted in 1,456 pRFs in the final grid.

#### Texture statistics features

As the first step of our voxelwise encoding model framework, we extracted a set of image-computable texture features which are meant to capture various aspects of local image structure (Figure 1 and Figure 2). We utilized the aforementioned P-S model (Portilla & Simoncelli, 2000), which has previously been shown to predict both human behavioral judgments of textures and neural responses in mid-level visual regions such as V2 and V4 (Freeman & Simoncelli, 2011; Freeman et al., 2013; Okazawa et al., 2015; Portilla & Simoncelli, 2000). Full details of the model construction and its motivation are given in (Portilla & Simoncelli, 2000) and (Freeman & Simoncelli, 2011). Here, we provide a brief description of the model components and describe how we have incorporated it into our pRF modeling framework.

The first stage of the model consists of using a steerable pyramid (Simoncelli & Freeman, 1995) to decompose each image into a set of orientation and frequency sub-bands (we used 4 orientations and 4 frequency bands). This step was implemented using the Python package *pyrtools* to construct a steerable pyramid in the frequency domain. Before processing images, we converted images to grayscale using the ITU-R BT.709-2 standard, which consists of multiplying the R,G,B channels by [0.2126, 0.7152, 0.0722] and taking their sum, and resampled images to a resolution of 240 × 240 pixels using bilinear resampling. The steerable pyramid results in spatial maps of complex-valued coefficients at each scale and orientation, from which we can compute the real part and the magnitude, which correspond approximately to the responses of V1 simple and complex cells, respectively (Freeman & Simoncelli, 2011). The four frequency bands output by our pyramid were centered at approximately 0.9, 1.8, 3.6, and 7.1 cycles per degree (cpd), and the four orientation bands were centered at 0°, 45°, 90° and 135°. The steerable pyramid additionally computes high-pass and low-pass residual images, as well as a partially-reconstructed low-pass image representation at each scale.

Our complete texture statistics model included 10 total subsets of features, resulting in 641 total features (see Table 1-1), each of which we computed at each pRF grid position. For simplicity, we have divided the features into two sub-groups for several of our analyses (e.g., Figure 5). The first sub-group, which we have termed “lower-level texture features”, consists of marginal statistics (such as mean and variance) computed from either the raw image luminance values, or from the outputs of the steerable pyramid. The second set of features, which we have termed “higher-level texture features”, consists of higher-order correlations computed from the steerable pyramid, generated by either correlating different orientation/scale channels of the pyramid, or spatially shifted versions of the same channel of the pyramid. By virtue of these cross-correlations and auto-correlations, the “higher-order” features are able to capture a higher degree of complexity than the “lower-level” model features, exhibiting sensitivity to properties like periodic, spatially repeating structure, and junctions made by contours of different orientations (compare top 4 and bottom 6 panels in Figure 2). When computing each of these features, we used the Gaussian profile for the pRF of interest (Equation (1)) as a weighting matrix (similar to the “pooling region” used in Freeman and Simoncelli, 2011). All feature extraction steps after the initial steerable pyramid computation were done using custom code in PyTorch.

**Figure 4:**
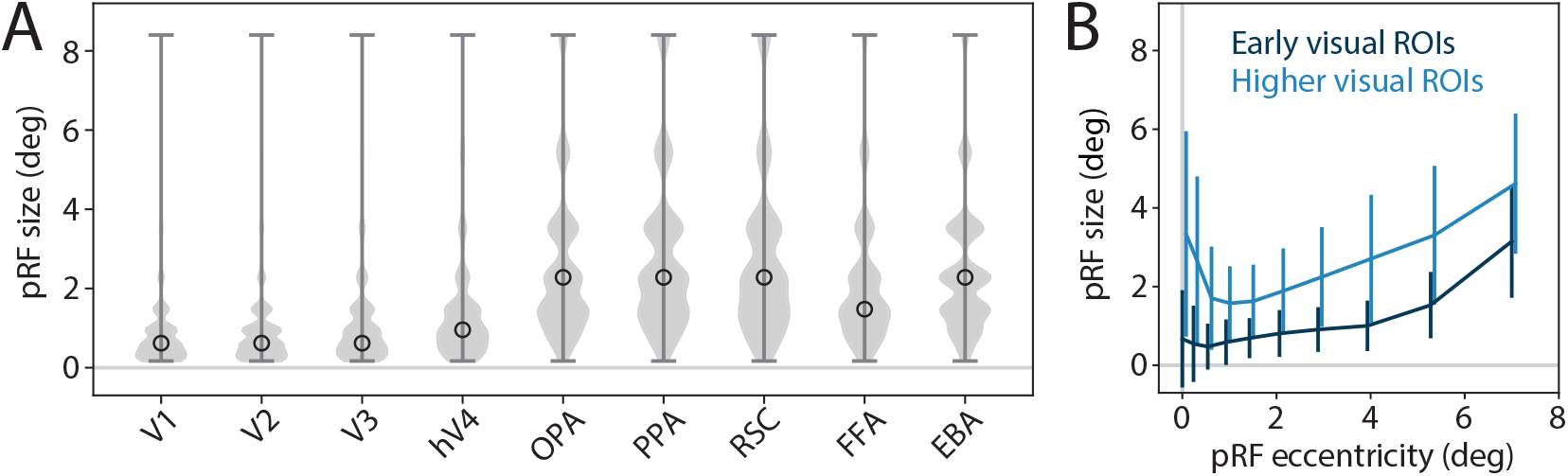
Population receptive field (pRF) parameters estimated using the texture statistics encoding model. See *Methods: Texture statistics encoding model* for details on pRF grid and fitting procedure. **(A)** Distribution of pRF size (*σ*) across voxels in each ROI. Distributions include voxels pooled across all subjects, open circle indicates the median. **(B)** Relationship between pRF *σ* and eccentricity, plotted for voxels pooled across early visual areas (V1, V2, V3, hV4; dark blue) or higher visual areas (OPA, PPA, RSC, FFA, EBA; light blue), across all subjects. Values of *σ* are averaged over all voxels having the same preferred eccentricity; error bars reflect the mean *±* standard deviation across *σ* values within an eccentricity.

**Figure 5:**
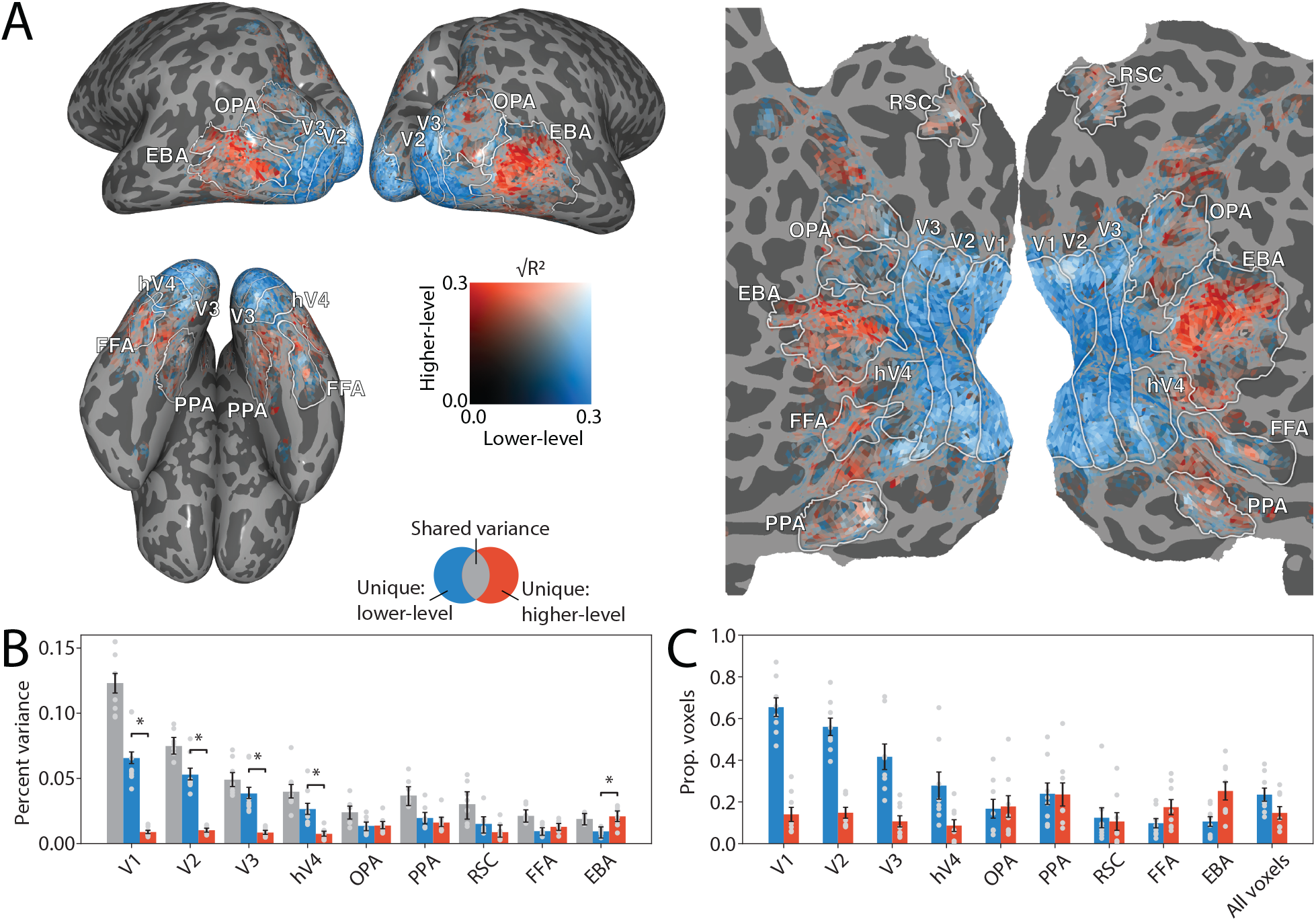
The unique variance explained by the higher-level texture features increases from lower to higher visual areas. Unique variance explained by the lower-level and higher-level features was measured using a variance partitioning analysis, see *Methods: Variance partitioning*. **(A)** Percent variance (units of 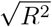) uniquely explained by the lower-level (shades of blue) and higher-level (shades of red) features, plotted on an inflated cortical surface from three viewpoints (left) or a flattened cortical surface (right) for one example subject, S1 (see Figure 5-1 for all subjects). Transparent voxels indicate that little variance was uniquely attributable to either the lower- or higher-level features, while white voxels indicate a large proportion of unique variance for both feature types. **(B)** Summary of the variance partitioning results averaged across ROIs and subjects. Gray bars indicate variance shared between the lower-level and higher-level features, while blue and red bars indicate variance unique to each set of features. Bar heights and errorbars indicate median and confidence intervals (99%) for the average unique and shared variance, obtained by bootstrapping the images when computing *R*^2^ (see *Methods: Variance partitioning* for details). All confidence intervals are significantly greater than zero (one-tailed *p*-values computed using a bootstrap test; corrected for multiple comparisons; *q* =0.01). Asterisks above pairs of bars indicate a significant difference between the lower-level and higher-level features (two-tailed *p*-values computed using a bootstrap test; corrected for multiple comparisons; *q* =0.01). Light gray dots indicate the mean variance explained for each individual subject. See Table 5-1 for the significance of individual subjects. **(C)** Proportion of individual voxels in each ROI that had a significant amount of variance uniquely explained by the lower-level (blue bars) and higher-level (red bars) texture features (one-tailed *p*-values computed using a bootstrap test; corrected for multiple comparisons; *q* =0.01; see *Methods: Variance partitioning* for details). Gray dots indicate individual subjects, bar heights and error bars indicate mean *±*1 SEM across subjects. The rightmost bars (“All voxels”) indicate the proportion of voxels that were significant across the entire analyzed visual cortex region, without regard to ROI definitions.

The first set of lower-level features, termed *pixel* features (6 features), consists of the pRF-weighted minimum, maximum, mean, variance, skew, and kurtosis of the raw pixel luminance values. The second and third sets of features, *energy-mean* (16 features) and *linear-mean* (16 features) features, are the pRF-weighted mean of the magnitude and real part, respectively, of each steerable pyramid feature channel (4 orientations x 4 scales). The fourth set of lower-level features, *marginal* features (11 features), includes the pRF-weighted skew and kurtosis of the low-pass pyramid reconstruction at each scale, and the pRF-weighted variance of the high-pass residual image.

The first set of higher-level texture features consists of autocorrelations of the steerable pyramid features, which are computed by correlating each feature map with spatially-shifted versions of itself – allowing these features to capture the repetition of similar elements across spatial positions. Autocorrelations were computed from the magnitude of each main pyramid feature channel (*energy-auto*; 272 features), the low-pass reconstruction at each scale, and the high-pass residual image. The autocorrelations of the low-pass reconstructions and the high-pass residual image were treated as a single combined group of features for the variance partitioning analysis (*linear-auto*; 98 features), but were treated separately when performing PCA (see below), as this yielded higher overall model accuracy. To compute each autocorrelation matrix, we cropped out a square region of the image that spanned approximately *±* 2*σ* from the pRF center. We then computed the weighted 2-dimensional autocorrelation over this cropped image region, using the pRF profile (cropped in the same way) as a weighting matrix. From this matrix, we retained a fixed number of pixels from the center. The number of pixels (i.e. spatial shifts) retained was adjusted based on the scale of the feature map under consideration; the total number of pixels retained ranged from 3 pixels (shifts of *±* 1 pixel) for the lowest frequency maps, to 7 pixels (shifts of up to *±* 3 pixels) for the highest frequency maps. Since the autocorrelation matrix is diagonally symmetric, we retained only the unique values. The total number of autocorrelation features returned was independent of pRF size.

The remaining higher-level texture features consist of cross-correlations, computed by correlating different feature maps output by the steerable pyramid. All cross-correlations were computed using the entire image, weighted by the pRF profile. The first two subsets of cross-correlation features, *energy-cross-orient* (24 features) and *linear-cross-orient* (34 features) are cross-correlations of feature maps (either magnitudes or real parts) at the same scale, but with different orientations; these features can thus capture image elements that include multiple orientations, such as crosses and curved lines. The next subsets of features (*energy-cross-scale*; 48 features) were computed by correlating the magnitude of feature maps having the same orientation but different scales, after up-sampling the resolution of the map at the coarser scale and doubling its phase. These cross-scale comparisons are able to capture distinctions between different types of oriented elements in the image, such as object edges versus lines versus gradients. An additional subset of features (*linear-cross-scale*; 116 features) were computed similarly, but using the real or imaginary component of the feature maps only. An additional two subsets of features were computed by correlating the low-pass residual image with spatially shifted versions of itself (within-scale) or with the lowest frequency pyramid feature map (cross-scale). The within-scale group of these features were included in the *linear-cross-orient* group for the variance partition analysis, while the cross-scale group were included in the *linear-cross-scale* group for the variance partition analysis, but these groups were treated separately when performing PCA (see next paragraph), as this grouping tended to result in better model performance.

To reduce the dimensionality of the texture model feature space and prevent over-fitting, we performed principal components analysis (PCA) on the higher-level texture features before using them to construct encoding models. PCA was performed within each pRF separately, within each subset of higher-level texture features individually (note that some feature groups were further subdivided when performing PCA, see above paragraphs). No dimensionality reduction was performed on the lower-level texture features. Where PCA was used, the principal components were always computed using the training data only, for one subject at a time, and all data for that subject including the validation data was projected into the same subspace. We retained the minimum number of components necessary to explain 95% of the variance. Since PCA was performed on the features from one pRF at a time, this meant that the dimensionality of the features was not required to be the same across all pRFs.

#### Model fitting procedure

To construct the texture statistics encoding model, we modeled each voxel response as a weighted sum of the texture statistics features corresponding to each image and each pRF (plus an intercept). As described in the previous section (*Methods: Texture statistics features*), texture statistics features were computed in a spatially specific manner, such that the feature activations for a given image depend on the pRF parameters *x*_*0*_, *y*_*0*_ and *σ*. We solved for the weights of the encoding model for each voxel using ridge regression (L2-regularization), as used in previous work (Güçlü & van Gerven, 2014; Huth et al., 2016; Wehbe et al., 2014). To improve the regression fits, we used a banded ridge regression method (Nunez-Elizalde et al., 2019) which incorporated two different ridge regularization (*λ*) parameters, one corresponding to the lower-level texture features and one corresponding to the higher-level texture features. Both *λ* parameters were selected on a per-voxel basis from a set of 10 candidate *λ* values logarithmically spaced between 0 - 10^5^. To sample all combinations of two *λ* values, we created a grid of all 100 possible *λ* combinations. Cross-validation was used to determine the best *λ* parameters for each voxel; the full cross-validation procedure is as follows (see also Figure 1).

First, we held out a validation set of *∼*1,000 images from the *∼*10,000 total images for each subject. This set of images always consisted of the shared images in which every subject saw the same images (note that for subjects who did not complete the entire NSD experiment, there were fewer images in this validation set; minimum was 907 images; see *Methods: Acquisition and preprocessing of fMRI data*). The remaining *∼*9,000 images made up the training data. Following this, we held out a random 10% of the training data as a nested validation set. This nested set was used to select the ridge parameter, as well as the best pRF parameters for each voxel. Using the remaining 90% of the training data, we then estimated regression weights for each of our candidate *λ* values, as well as for each of our candidate pRF models. Based on the estimated weights, we computed the loss for each lambda and each pRF, by generating a prediction of the nested validation data and computing the sum of the squared error for this prediction. We then selected the best pRF parameters and *λ* values for each voxel based on which values resulted in the lowest loss. The resulting pRF parameters and regression weights made up the final encoding model for the voxel. Finally, to estimate overall model accuracy, we generated predicted responses of each voxel on the held-out validation set and computed the coefficient of determination (*R*^2^) between the actual and predicted response. To make plots of *R*^2^ in surface space for each subject (i.e., Figure 3A), we utilized PyCortex (Gao et al., 2015).

#### Permutation testing

For each voxel, we used a permutation test to evaluate whether the texture statistics encoding model resulted in higher than chance accuracy at predicting the validation set data. To perform the permutation test, we randomly shuffled (1,000 times) the image labels corresponding to each voxel activation pattern (voxel activations were already averaged over presentations of each repeated image before this shuffling was performed; see *Methods: Acquisition and preprocessing of fMRI data*). Shuffling was always performed within the training set, validation set, and nested validation set (see *Methods: Model fitting procedure*) separately. For each shuffling iteration, we performed the regression procedure from scratch, including fitting the model weights and computing *R*^2^. However we did not re-fit each voxel’s pRF on the shuffled data, instead we used the best pRF for each voxel as determined from the intact data, and only re-fit the regression weights. To compute a one-tailed *p*-value for each individual voxel, we calculated the number of iterations on which the shuffled *R*^2^ was *≥*the real *R*^2^, divided by the number of iterations. We then performed FDR correction on the *p*-values for all voxels from each subject (*q*=0.01; Benjamini and Hochberg, 1995). We used the result of this significance test as a mask to determine which voxels to include in the subsequent variance partitioning analyses.

#### Variance partitioning

In order to determine the unique contribution of different subsets of texture features to encoding model predictive accuracy, we used a variance partitioning analysis (see Groen et al., 2012; Lescroart and Gallant, 2019 for similar approaches). When performing this analysis, we always restricted the set of voxels to those that had above-chance accuracy for the full texture statistics encoding model (see previous section). The overall approach was as follows. First, we fit the full texture statistics encoding model with all feature subsets concatenated, using the method described in the previous section. We computed the prediction accuracy of the concatenated model, 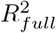. Then, we progressively removed one set of features at a time from the full model, and fit the weights of each partial model. When constructing the partial models, we always used the features corresponding to the voxel’s best pRF (as determined based on fitting the full concatenated feature space). This ensured that the differences between models were due solely to differences in features and not to changes in the estimated pRF parameters. Then, for each partial model, we generated a prediction of the validation set data and computed *R*^2^ for the partial model,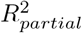. Finally the unique variance attributable to each feature subset was computed as:

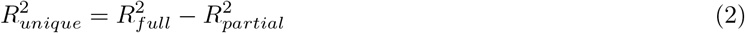

Where 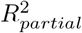 denotes the partial model where the feature set of interest was removed. We performed two versions of the variance partitioning analysis: in the first version, we grouped all of the lower-level or all of the higher-level texture features together (Figure 5); for the second version, we analyzed each of the 10 feature subsets individually (Figure 7, Figure 8). For the first analysis, we additionally report the variance that is shared between the lower- and higher-level feature subsets. The shared variance between two feature subsets *A* and *B*, was computed as:

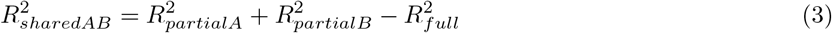

To evaluate the significance of the unique variance explained by each feature subset, we used a bootstrapping analysis consisting of resampling with replacement. To make this analysis computationally feasible, we performed bootstrapping only on the validation set data (i.e., when computing *R*^2^ but not when fitting model weights). We performed 1,000 iterations of the bootstrapping test; on each iteration we resampled with replacement *n* images from the total *n* validation set images (where *n* is typically 1,000, but could be fewer for the subjects who did not complete all sessions; see *Methods: Acquisition and preprocessing of fMRI data*). We used the same resampling order for the image labels and for the voxel data, so that the correspondence between images and voxel responses was intact, but the exact set of images included differed on each bootstrap iteration. Using the same resampling order, we computed *R*^2^ for the full model and *R*^2^ for each partial model, then used these to compute 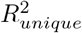 for each feature set, as described above. This resulted in a distribution of 1,000 values for 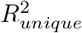. We then used this distribution to compute a *p*-value for whether 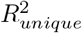 was significantly higher than zero, by computing the number of iterations on which 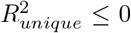 and dividing by the number of iterations. The *p*-values for each subject were FDR corrected across all voxels (*q*=0.01; Benjamini and Hochberg, 1995).

**Figure 6:**
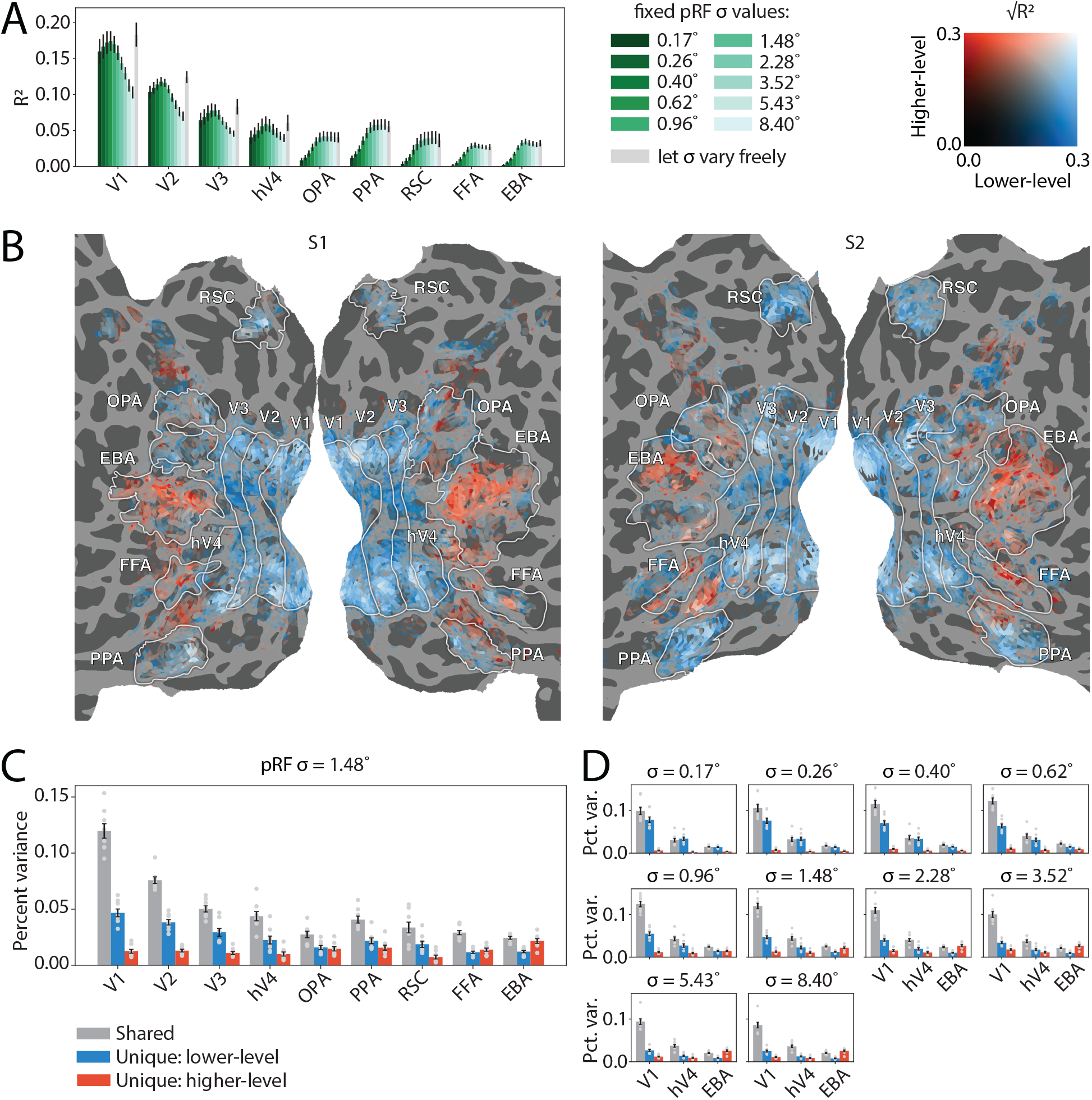
The difference in feature sensitivity between early and higher visual areas is not dependent on differences in receptive field size. We re-fit the entire texture statistics encoding model with a single fixed pRF size (*σ*=1.48 degrees) for all voxels, see *Methods: Fitting with fixed pRF size parameter* for details. **(A)** Overall accuracy (*R*^2^) of models in which pRF size (*σ*) was fixed at a single value for all voxels, averaged across voxels in each ROI. Each colored bar represents a different *σ* value; gray bar represents the model in which sigma was allowed to vary across voxels (similar to Figure 3B). Bar height and errorbars indicate the mean *±* SEM across subjects. **(B)** Percent variance (units of 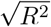) unique to the lower-level (shades of blue) and higher-level (shades of red) texture features, shown on a flattened cortical surface for two example subjects. **(C)** The percent of variance that was shared among feature types (gray bars), unique to the lower-level texture features (blue bars), or unique to the higher-level texture features (red bars), averaged over voxels within each ROI. Bar heights and error bars indicate mean *±*1 SEM across subjects, light gray dots indicate individual subjects. **(D)** Variance partitioning analysis performed for other values of *σ*, shown for selected ROIs V1, hV4, and EBA. Bar heights and error bars are as in **(C)**.

**Figure 7:**
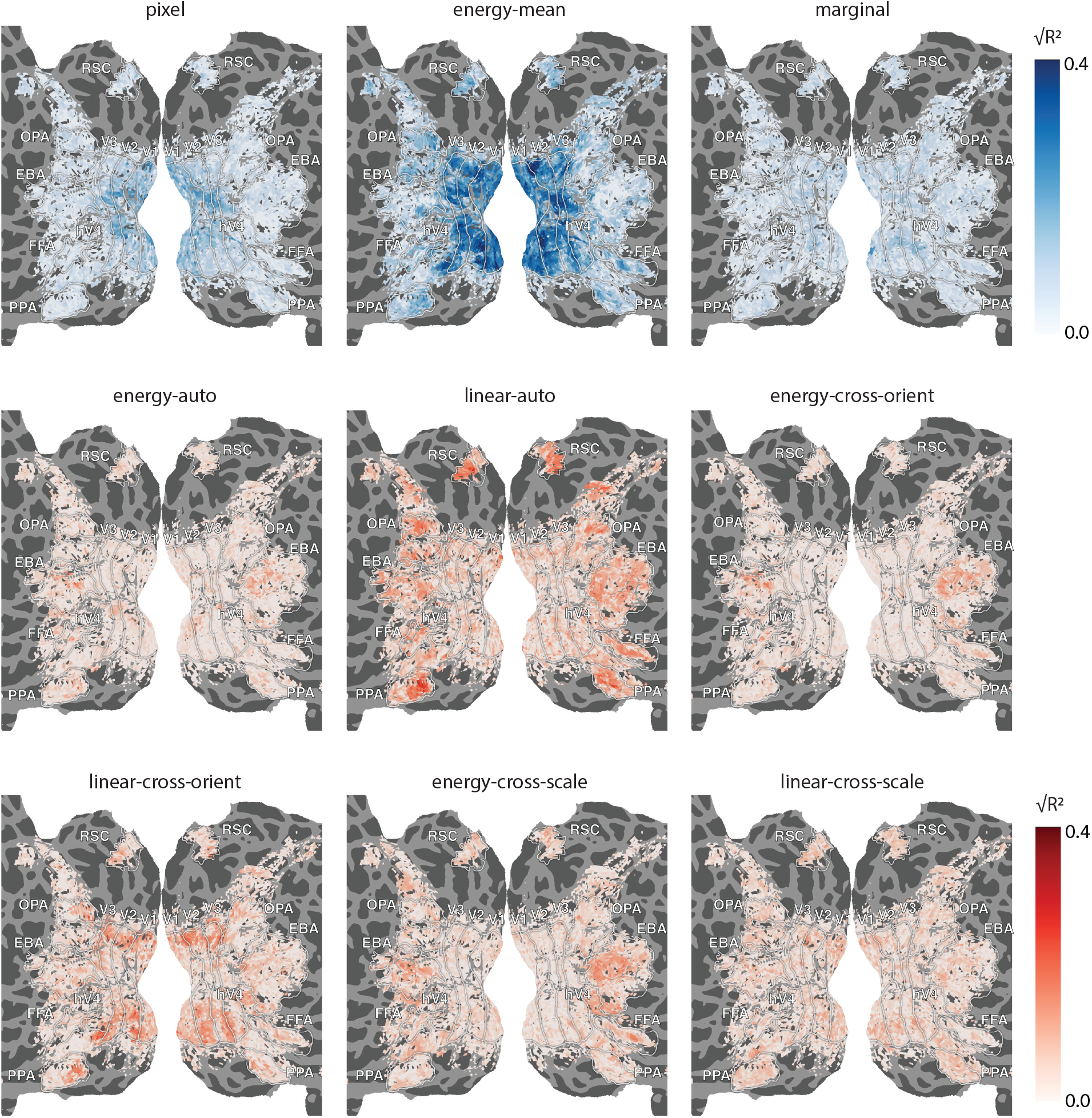
The organization of neural populations sensitive to each subset of texture model features, shown for one representative NSD subject (S1). See Figure 7-1 for the full set of 8 subjects. Each panel shows thepercentage of encoding model variance (units of 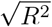) uniquely explained by each individual feature type, shown on a flattened cortical surface (see *Methods: Texture statistics features* for details on feature types). The maps in shades of blue (top row) correspond to subsets of the lower-level texture features, while maps in shades of red (bottom two rows) correspond to subsets of higher-level texture features. Note that the *linear-mean* features are not plotted here, as they explained little unique variance in any ROI (see Figure 8).

**Figure 8:**
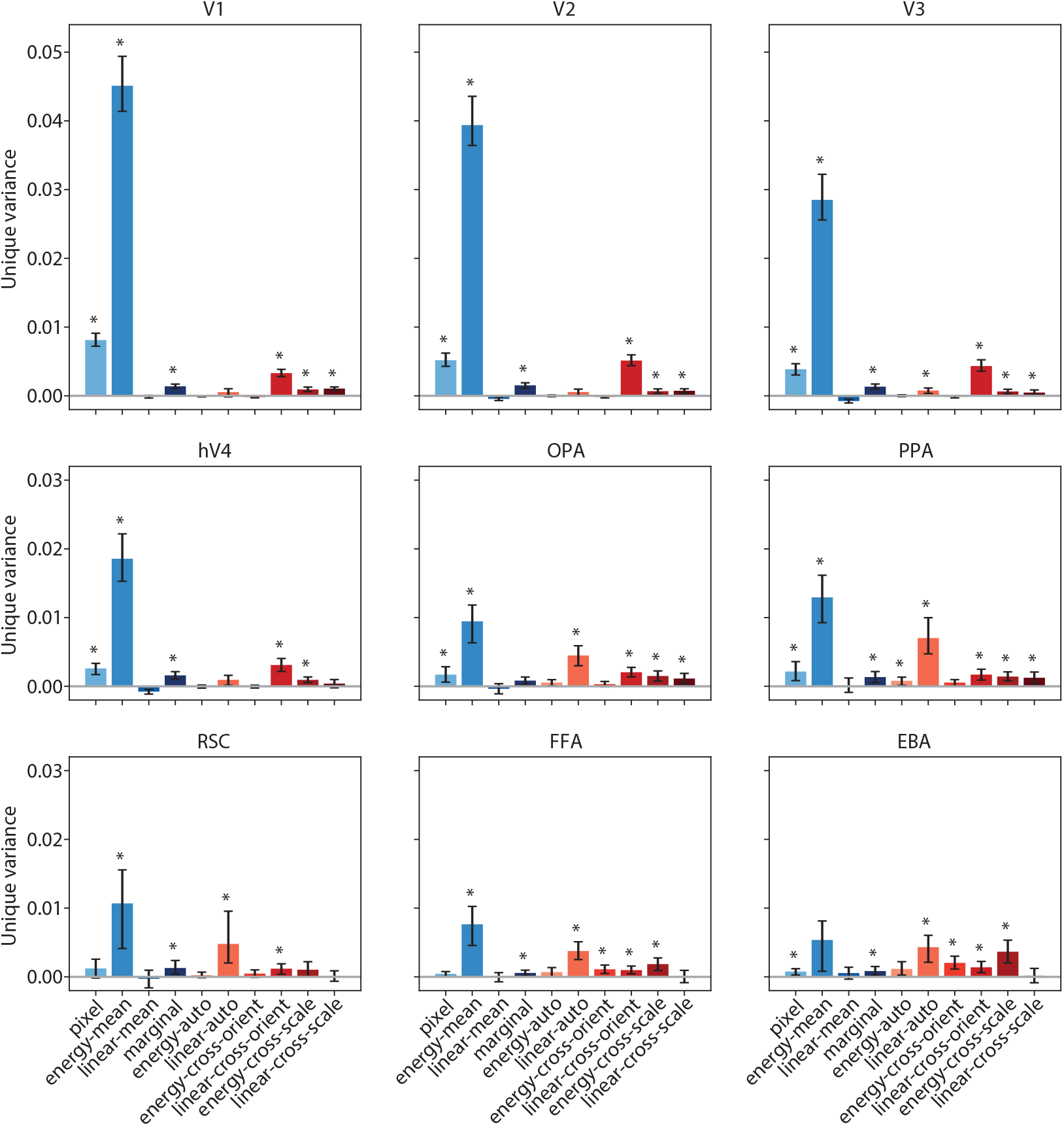
Results of variance partitioning analysis across all feature types, summarized at the ROI level. Each panel shows the proportion of model variance uniquely explained by each feature type, for one ROI, averaged across all subjects. Bar heights and errorbars indicate median and confidence intervals (99%) for the average unique variance, obtained by bootstrapping the images when computing *R*^2^. Asterisks (*) above each bar indicate that the unique variance was significantly greater than zero (one-tailed *p*-values computed using a bootstrap test; corrected for multiple comparisons; *q* =0.01; see *Methods: Variance partitioning* for details). Bars in shades of blue indicate feature subsets belonging to the “lower-level” group, while bars in shades of red indicate feature subsets belonging to the “higher-level” group. See Table 8-1 for the number of individual subjects in which unique variance was significant for each feature subset.

We additionally computed whether the unique variance for each feature set was significant at the ROI-averaged level for each subject. This was done by averaging the bootstrap distributions for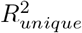 across all voxels in each ROI (always averaging values for the same bootstrapping iteration together), and then computing a one-tailed *p*-value using the same method described above. This resulted in a single value for each ROI, each subject, and each feature set. We then performed multiple comparisons correction across all values using the Holm-Bonferroni method (*q* =0.01; Holm, 1979). We used the Holm-Bonferroni method because the FDR procedure is not appropriate for controlling for a relatively small number of comparisons. The number of subjects with significant unique variance explained for each feature set is reported in Table 5-1 and Table 8-1. To test whether the average unique variance across all subjects was significant, we used the same averaging method just described but additionally averaged the bootstrap distributions over subjects before computing *p*-values, and corrected for multiple comparisons as described above.

To compute whether the unique variance explained by the lower- and higher-level texture features was significantly different for each ROI, we used the ROI-averaged bootstrap distributions to compute the distribution of differences between the lower-level unique variance and the higher-level unique variance. We then computed a two-tailed *p*-value by computing the proportion of iterations for which the difference was positive and the proportion for which the difference was negative, taking the minimum and multiplying by 2. We again performed multiple comparisons correction using the Holm-Bonferroni method (*q* =0.01).

#### Fitting with fixed pRF size parameter

To dissociate the effects of pRF size and feature selectivity, we performed an additional analyis in which the pRF size parameter (*σ*) was fixed at a single value for all voxels. To achieve this, we performed our entire model fitting pipeline from scratch (see *Methods: Model fitting procedure*) with a restricted grid of candidate pRFs. This restricted pRF grid consisted of all the pRFs in our main grid having the *σ* value of interest. Thus, the voxels could be fit with different pRF centers, but had to have the same pRF size. We performed this entire procedure for each of the 10 *σ* values in our original pRF grid.

### Alternative low-level visual models

To assess how the texture statistics model’s ability to predict neural responses compares to that of other models, we implemented several commonly-used models of low-level visual features: a GIST model (Oliva and Torralba, 2001), a complex Gabor model (Henderson et al., 2022; St-Yves and Naselaris, 2017), and the first two layers of a pretrained Alexnet convolutional neural network model (Krizhevsky, 2014). The GIST model consists of spectral features (orientation and spatial frequency) that are coarsely localized in space, and was implemented using Matlab code provided by the authors of Oliva and Torralba, 2001. We evaluated the GIST model with two levels of spatial resolution: a 2×2 grid and a 4×4 grid. To make the GIST model as comparable as possible to our texture statistics model, we used 4 orientation channels and 4 frequency channels, which matches the number of channels included in the steerable pyramid. Similarly, when implementing the Gabor model, we also used 4 orientations and 4 spatial frequencies (0.36, 1.03, 2.97, and 8.57 cycles/deg). Each Gabor model feature was computed by filtering the image with two sinusoids that were 90° out of phase, squaring the output of these two filters, summing the two outputs, and taking the square root (see Henderson et al., 2022 for details on construction of a similar model). To extract features from the Alexnet model, we used the pretrained model weights for Alexnet available from the PyTorch model zoo, and we extracted activations from layers Conv1 and Conv2 (following the rectifying nonlinear activation function). More details on Alexnet can be found in (Krizhevsky, 2014); briefly, it is a convolutional neural network model trained to perform 1000-way object classification. Thus in contrast to the Gabor, GIST, and texture statistics models, which each consist of hand-designed feature weights, the Alexnet model is task-optimized and may include object-specific features.

For both the Gabor and Alexnet models, we incorporated spatial pRF parameters into the construction of the model, similar to how the pRF was incorporated into the texture statistics model. Essentially, this procedure consists of extracting features in each pRF of the grid by taking a dot product of the relevant feature maps with each pRF (for similar approaches, see: Henderson et al., 2022; St-Yves and Naselaris, 2017). To simplify the fitting procedure, as well as making the models more comparable to the texture statistics encoding model, we used the same pRF parameters that had already been estimated using the texture statistics encoding model (see *Methods: Model fitting procedure*). This meant that when fitting the Gabor and Alexnet models, we only had to fit the feature weights (using the set of features that had been extracted from each voxel’s best pRF). For the GIST model, the pRF parameters were not incorporated into the fitting procedure; instead the GIST features included information from the entire image.

### Code Availability

All code required to reproduce our analyses is available at: https://github.com/mmhenderson/texturemodel.

## Results

To model neural selectivity for mid-level image structure, we constructed a forward encoding model based on image-computable texture statistics features (Figure 1 and Figure 2; see also *Methods: Texture statistics encoding model*). The texture statistics encoding model includes parameters that capture the spatial selectivity of voxels (population receptive fields or pRFs; Dumoulin and Wandell, 2008; St-Yves and Naselaris, 2017) as well as their selectivity for a range of features related to local image structure (computed using P-S statistics; Portilla and Simoncelli, 2000; described in more detail below). All parameters of the model were fit on a voxelwise basis, and overall accuracy for each voxel was quantified by generating predicted voxel responses to a set of held-out images that were not used during model training. We then computed the coefficient of determination (*R*^2^) for each voxel. To facilitate comparisons across visual regions with different functional roles in mid-level visual processing, we fit and evaluated the model across a large portion of occipotemporal cortex, as well as computing summary statistics for several functional ROIs: early retinotopic areas V1-hV4, scene-selective areas OPA, PPA, RSC, face-selective area FFA, and body-selective area EBA (see *Methods: Defining regions of interest (ROIs)* for details).

Our model includes a range features related to both low and mid-level properties of images. Lower-level properties include the activation in different orientation and frequency channels as extracted by a steerable pyramid (Simoncelli & Freeman, 1995), and the marginal statistics of the pixel luminances; we refer to these collectively as “lower-level” texture features (see *Methods: Texture statistics features* for more details). The model also includes features that capture higher-order (i.e., mid-level) structure in the image, including the cross-correlations between different orientation and frequency maps output by the steerable pyramid, as well as auto-correlations computed from individual steerable pyramid maps; we refer to these collectively as “higher-level” texture features. Figure 2 provides examples of image patches that result in high and low values for the first two principal components of each subset of model features, giving some intuition for the properties that are captured by each set of features. The top four panels illustrate the four subsets of lower-level model features, which capture relatively simple aspects of image structure such as mean luminance (*pixel* features; PC2) and the strength of horizontal and vertical orientations (*energy-mean* features; PC2). In contrast, the bottom six panels illustrate the types of properties captured by the higher-level model features, which are relatively more complex. For example, the *linear-auto* features appear to differentiate image patches according to the presence of high-frequency spatially repeating structure in the images (items like window blinds and zebra stripes), while PC1 of the *linear-cross-scale* features differentiates patches including white lines on a black background from patches including black lines on a white background. Other subsets of the higher-level model features, such as the *energy-cross-orient* features, appear to be related to conjunctions between differently oriented elements in the image (bent knees and elbows on a person, angles created by parts of an airplane), while PC1 of the *energy-cross-scale* features appears to be related to spatial frequency, but maintains some invariance to orientation. Some of the higher-level model features also appear to capture distinctions between image patches that are more angular or rectilinear versus those that include more organic and curvy shapes (see PC2 of the *energy-cross-orient* and *energy-cross-scale* features). While not all of the principal components are easy to describe in words, this illustration provides some intuition as to how our model can differentiate complex natural scene images according to multiple aspects of their mid-level image structure.

Overall, the texture statistics encoding model achieved good predictive accuracy (*R*^2^) across multiple visual areas, with particularly strong performance in early visual cortex (Figure 3A and Figure 3B, blue bars and Figure 3-1). Although performance of the model progressively declined from V1 through more anterior category-selective visual regions, performance remained moderately high in voxels throughout the visual hierarchy, with the majority of voxels in all visual ROIs having above-chance validation set accuracy (Figure 3D; one-tailed *p*-values obtained using permutation test, corrected for multiple comparisons; *q*=0.01). In addition, examining the spatial fit parameters of the model (Figure 4) demonstrates that the model recovers single-voxel pRFs with properties that are consistent with past work (Dumoulin & Wandell, 2008), namely the tendency of pRF size (*σ*) to increase from early to higher-level visual ROIs and the tendency of pRF size to scale with eccentricity. These results provide validation of our modeling framework and indicate that the model is able to capture a substantial portion of the response variance in voxels across multiple stages of the visual hierarchy.

We compared our model’s performance to several commonly used models for early vision, including the GIST model (implemented with a coarse 2×2 and a finer 4×4 spatial grid), a Gabor model, and the first two layers of a convolutional neural network, Alexnet (for details, see *Methods: Alternative low-level visual models*). The GIST and Gabor models were selected for comparison because they both capture spectral information, but not higher-order correlations, meaning they should have some feature overlap with the lower-level texture model features, but not the higher-level features. In contrast, Alexnet is a larger model trained on object classification, and thus may encode features not captured within the texture statistics model. In particular, the increase in feature complexity from Conv1 to Conv2 of AlexNet may allow it to capture higher-order correlation structure from images, similar to the higher-level texture features in our model. As shown in Figure 3B, our texture statistics model had comparable prediction accuracy to these other models, with texture model accuracy slightly exceeding that of the 2×2 GIST model, the Gabor model, and the first convolutional layer of Alexnet (see also Figure 3C). The texture model’s accuracy was exceeded slightly by the second convolutional layer of Alexnet and (in some higher visual areas) the 4×4 GIST model. In the case of Alexnet Conv2, the likely reason for this advantage is that Alexnet is able to capture a wider range of features than the texture statistics model because it has many more parameters and is task-optimized, as opposed to being based on hand-designed features. In the case of GIST, the difference was unexpected because the GIST features are based on Gabor filter outputs and are thus relatively low-level. On reflection, one possible explanation is that GIST includes spectral features extracted from multiple positions in the visual field, so it is capable of capturing higher-order aspects of response selectivity. For example, GIST can capture the responses of a voxel that is sensitive to vertical orientations in the upper left visual field and horizontal orientations in the lower right visual field. Thus, GIST is able to approximate some of the same features captured by higher-level texture features in our model - which may explain its slight advantage in higher visual areas with larger RF sizes (e.g., PPA). It is important to note that the main goal of our modeling was *interpretability*, rather than maximizing model accuracy or outperforming pre-existing models. That is, we developed a model that can be used to isolate the contributions of different mid-level texture features, a property that is not afforded by any of our comparison models. The fact that the texture model yields competitive performance with similar models provides a good indication that its features are reliably predictive of neural responses.

Given the high overall accuracy of the texture statistics encoding model, we next asked which features were most critical to its predictive performance. To test this, we performed a variance partitioning analysis (see *Methods: Variance partitioning*), where we sub-divided the model features into lower-level and higher-level texture features (see Figure 1), and determined both the percentage of variance that was uniquely attributable to each set of features and the percentage that was shared among the two sets (Figure 5). This analysis revealed key differences among visual areas. First, visualizing the unique variance values on a flattened cortical surface (Figure 5A and Figure 5-1) revealed a gradient from lower to higher visual areas, where voxels in the most posterior portion of occipital cortex tended to have more unique variance explained by the lower-level features (shades of blue), while more anterior voxels, particularly on the lateral surface of the brain, had progressively more variance uniquely explained by the higher-level texture features (shades of red). Averaging the unique variance values across voxels within each ROI further underscored this dissociation between areas: in early retinotopic areas V1-hV4, the lower-level texture features explained more unique variance than did the higher-level texture features, but in higher category-selective visual areas FFA and EBA, this trend reversed, with the higher-level features accounting for more unique variance on average (Figure 5B). Place-selective ROIs OPA, PPA, and RSC showed a more intermediate pattern, with the lower- and higher-level features explaining similar amounts of unique variance. Consistent with these patterns, a bootstrap significance test revealed that the average unique variance explained was significantly higher for the lower-level features than the higher-level features in V1, V2, V3, and hV4, but it was significantly higher for the higher-level features than the lower-level features in EBA (Figure 5B; two-tailed *p*-values computed with a bootstrap test; corrected for multiple comparisons; *q* =0.01; see *Methods: Variance partitioning*). In addition to significance at the ROI-averaged level, these differences were significant in all individual subjects in V1-V3, in 7 individual subjects in hV4, and in 4 individual subjects in EBA (Table 5-1). Furthermore, more individual voxels in early visual areas had significant unique variance for the lower-level features than for the higher-level features, while more voxels in FFA and EBA had significant unique variance for the higher-level features (Figure 5C; one-tailed *p*-values for single voxels computed using bootstrap test; corrected for multiple comparisons; *q* =0.01). Another trend evident in this analysis was that across all visual areas, a substantial portion of the variance was shared between the lower-level and higher-level texture features (gray bars in Figure 5B), suggesting some degree of feature overlap between the lower- and higher-level feature spaces.

The distinction between low- and high-level visual areas in the previous analysis is consistent with the interpretation that these areas represent features at different levels of complexity, with early areas representing more low-level aspects of image structure and higher visual areas representing higher-order statistics. However, another factor that could potentially contribute to this distinction is receptive field size. In our procedure for computing texture statistics features, the pRF size parameter (*σ*) determines the size of the spatial weighting function used when computing each texture feature (see *Methods: Texture statistics features* for details). This implies that for larger *σ* values, the higher-order texture statistics features may be more informative, since they incorporate information about a larger portion of the visual field. Since voxels in higher visual cortex tend to have larger *σ* than voxels in early visual cortex (see Figure 4A), it is possible that the ability of higher-level texture features to explain more variance than lower-level features within higher visual areas is driven only by larger receptive field sizes. To evaluate this possibility, we constructed new versions of our encoding models in which *σ* was fixed at a single value, and then re-fit the entire encoding model with all voxels having the same *σ* (see *Methods: Fitting with fixed pRF size parameter* for details). Consistent with the difference in estimated receptive field size between areas, this procedure resulted in highest performance in early areas when the fixed *σ* value was relatively low, but highest performance in higher areas when *σ* was relatively high (Figure 6A). To allow for a balanced comparison across early and higher visual areas, we selected a *σ* close to the middle of our range of candidate values, 1.48°, for further inspection.

Importantly, when *σ* was fixed at 1.48° we replicated the key findings of our original model (Figure 6B-C). We again found that in posterior early visual areas, the lower-level texture features explained a greater proportion of the model variance than the higher-level texture features, but in more anterior areas this pattern began to reverse, with the higher-level texture features explaining a progressively larger percentage of the variance. This pattern was observed across a range of *σ* values (Figure 6D), though not for the lowest *σ* values (note that as described previously, small *σ* values resulted in low overall *R*^2^ for higher visual areas). The observation that a distinction between the coding properties of low- and high-level visual areas can be recovered even when *σ* is fixed at a single value for all voxels indicates that the difference between these areas involves a true difference in feature selectivity, not only a difference in the scope of spatial selectivity. This result is also consistent with (Freeman et al., 2013), who found no correlation between the receptive field size of V1 and V2 neurons and their relative sensitivity to higher-level texture statistics.

The previous analyses suggest a dissociation between low and high-level visual areas in terms of which texture statistics features best explain their responses. However, it is not yet clear whether all subsets of the lower- and higher-level texture features contribute equally to model performance in a given brain region, or whether there is redundancy among the feature subsets. Thus, to more precisely determine which texture features were most uniquely predictive of neural activation, we next performed a second variance partitioning analysis where we considered each subset of both the lower-level and higher-level feature groups separately (a total of 10 feature types; see *Methods: Texture statistics features* for details).

As shown in Figure 7, Figure 8, and Figure 9, this analysis revealed that overall model accuracy reflected unique contributions from multiple feature subsets. Across all areas, the feature subset which uniquely explained the largest proportion of the texture model’s *R*^2^ was the *energy-mean* features, (Figure 7, top middle panel) which are analogous to a model of complex cell responses in V1. In early areas, a small amount of variance was also explained by the *pixel* features, which include marginal statistics computed from the image luminance histogram, and thus may capture low-level properties like overall image brightness or contrast (top left panel). A portion of the variance in some early areas was also explained by the *linear-cross-orient* features (bottom left panel), a higher-level feature subset which consists of cross-correlations between the real parts of steerable pyramid bands with different orientations. These features may capture higher-order structure generated from the combination of different orientations, such as corners, angles and other contour junctions (see Figure 2 for examples). Notably, the highest values of unique variance for the *linear-cross-orient* features tended to be observed for voxels sensitive to the vertical meridians in early visual cortex (i.e., boundaries between V1 and V2, and between V3 and hV4). Given the previously-reported relationship between pRF polar angle and preferred orientation (Freeman et al., 2011; Roth et al., 2022), this retinotopic relationship may suggest these features are related to a comparison between vertical and horizontal orientations, as well as having some association with visual field position. Moving into more anterior regions, other higher-level features began to explain a larger proportion of the variance. The *linear-auto* features, which are computed based on spatial autocorrelations, explained a moderate proportion of variance across several high-level visual areas, including voxels in and near the category-selective areas (center panel) – this feature subset may capture periodic, spatially repeating textural elements in the images (see Figure 2). Other feature subsets which explained a notable amount of unique variance in higher-level visual cortex included those computed from higher-order statistics of the steerable pyramid magnitude features, including the *energy-cross-scale* features, as well as the *energy-cross-orient* features.

**Figure 9:**
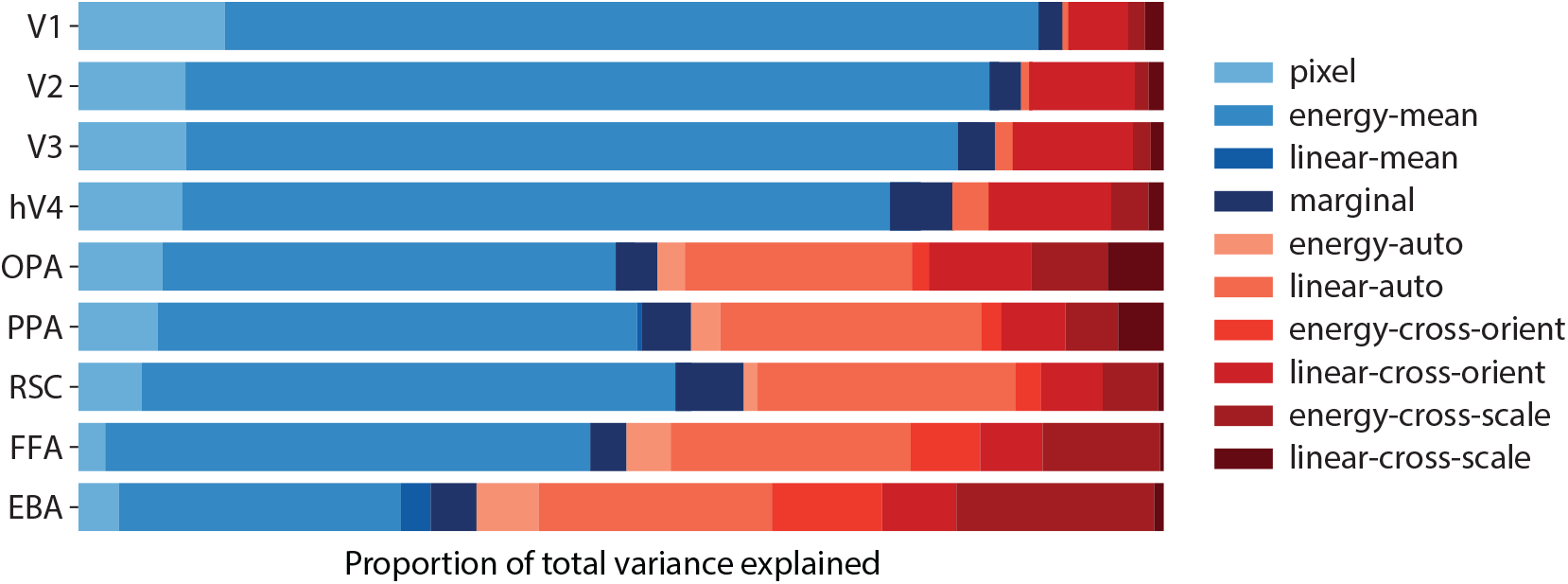
Proportion of the total model variance in each ROI explained by each feature type. Variance partitioning analysis is performed in the same way as in Figure 8, but each unique variance value is expressed as a proportion of the model’s total *R*^2^. Results are averaged across all subjects. Blue shades indicate lower-level model features, while red shades indicate higher-level model features. See *Methods: Variance partitioning* for more details.

Additionally, the spatial pattern of unique variance explained by each feature type suggested some heterogeneity within individual ROIs. For example, different portions of EBA appeared to be differentially predicted by the *linear-auto, energy-cross-scale*, and *energy-cross-orient* features, and higher-level features like the *linear-cross-orient* features explained a larger amount of variance in the posterior portion of PPA than the anterior portion. Some of this variability may be attributable to differences in the model’s overall accuracy across voxels within an ROI (Figure 3A), but it may also suggest that these ROIs contain sub-regions that can be delineated based on mid-level response properties.

To quantify these effects, we averaged the unique variance across voxels in each ROI (Figure 8, Figure 9). Again, this analysis demonstrated a primary role for the *energy-mean* features in all ROIs, with this feature subset dominating the most strongly in early visual areas. The unique variance explained by the *energy-mean* features was significant at the subject-averaged level in all ROIs except EBA, and was significant in all individual subjects in V1, V2, V3, and hV4 (Table 8-1; one-tailed *p*-values computed using bootstrap test; corrected for multiple comparisons; *q* =0.01; see *Methods: Variance partitioning* for details). Though the absolute amount of unique variance contributed by other feature subsets was modest in comparison to the *energy-mean* features, other feature subsets nonetheless contributed a significant amount of unique variance in each ROI. The *pixel* features made significant contributions in several areas, with highest unique variance contributed in V1 and V2 (all 8 subjects individually significant in V1 and V2). As suggested in the previous paragraph, the *linear-cross-orient* features also contributed to the variance explained in many areas, with unique variance explained significant at the subject-averaged level in all ROIs (significant in all 8 subjects in V1 and V2). Another higher-level feature subset that stood out was the *linear-auto* features, which explained a significant amount of unique variance in V3 as well as OPA, PPA, RSC, FFA and EBA. In general, the relative proportion of unique variance explained by these higher-level texture statistics features increased from early visual areas to higher visual areas (Figure 9). These results suggest that the performance of the texture statistics encoding model reflects contributions from multiple feature subsets within the model, with the relative importance of these feature subsets varying among visual areas. Areas across the visual cortex may encode complementary aspects of image texture, resulting in a population representation that spans a large portion of the full texture statistics feature space.

To further explore the representational space defined by the texture statistics model, we performed PCA on the unique variance values for each of the 10 feature subsets (Figure 10). PCA identified a set of weights in voxel space that project the unique variance values onto a lower-dimensional subspace. Visualizing these weights for the first four principal components reveals several large-scale organizational motifs across visual cortex. The first principal component (PC1) had its highest weights for voxels in early visual cortex, with a high score for the *energy-mean* features – this result is not surprising given the high magnitude of unique variance values for the *energy-mean* features in early visual cortex (Figure 7). The second principal component (PC2) had high positive weights for voxels in higher visual cortex and negative weights in early visual areas, especially for voxels sensitive to the central visual field. PC2 had a high positive score for the *linear-auto* features and negative score for the *pixel* features (Figure 10C). Plotting the top and bottom images for PC2 indicates a potential relationship with image scale, as well as the presence of human figures and/or actions. Interestingly, the third principal component (PC3) seemed to create a rough division within early visual cortex based on retinotopy, having positive weights for voxels sensitive to more central and horizontal visual field positions, and negative weights for those sensitive to vertical visual field positions. This is consistent with the observation that PC3 has a strong negative score for the *linear-cross-orient* features, which also had high unique variance values for the vertical-meridian-preferring populations in early visual cortex (Figure 7); as mentioned previously, these populations are also likely to be selective for vertical orientations. Consistent with this, PC3 tended to be most activated by scenes including a strong representation of the horizon (meaning voxels with a negative weight on PC3 were negatively associated with horizon-dominated scenes) – possibly reflecting sensitivity for the higher-order statistics associated with naturalistic outdoor scenes. Finally, PC4 appeared to be most positively weighted for voxels in face- and body-selective regions (FFA, EBA), and more negatively weighted in PPA and RSC (especially for S1) as well as in periphery-preferring regions of early visual cortex. PC4 had its highest positive score for the *energy-cross-scale* features, and a negative score for the *linear-cross-orient* features. Based on the top and bottom images for this component, PC4 appears to be sensitive to the difference between images including animals or humans versus scene images that were strongly geometric in appearance. This is consistent with past work suggesting that sensitivity to mid-level features associated with object animacy is a dominant axis of visual cortex organization (Konkle and Caramazza, 2013). Taken together, these results suggest that several large-scale organizational properties of visual cortex can be recovered based only on the patterns of unique variance across feature subsets in our model. Therefore, texture statistics features may provide an interpretable form of scaffolding for the emergence of higher-level semantic representational axes in the brain.

**Figure 10:**
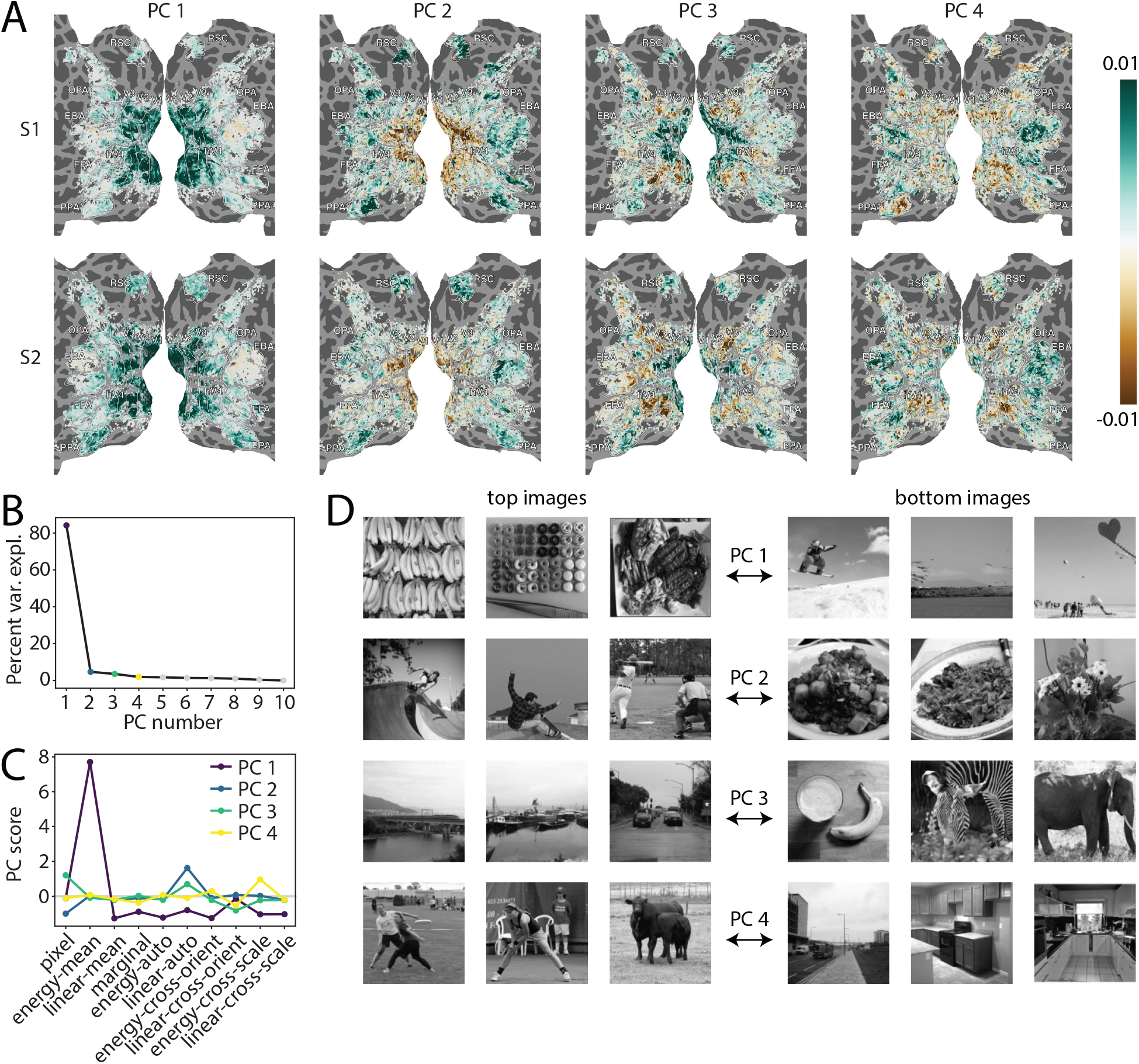
Principal components analysis (PCA) performed on the unique variance values for each subset of texture statistics features. We concatenated the unique variance values for each feature subset (i.e, data in Figure 7) across all subjects, and then performed PCA to learn a set of weights that projects the unique variance values onto a lower-dimensional subspace. **(A)** The weights (loadings) for the first four PCs are plotted on a flattened cortical mesh for two example subjects (top row=S1, bottom row=S2). Similar results were obtained for the other 6 subjects. **(B)** The percent of variance explained by each principal component. **(C)** The score for each of the feature subsets with respect to the first four principal components (each colored line indicates a different principal component). **(D)** The most activating (left) and least activating (right) images are shown for each of the first four principal components. To obtain these images, we identified the images that were overlapping across all subjects (907 images) and concatenated the voxel responses to these images across all subjects. PCA weights were then used to project these voxel response patterns into principal component space, and we identified the 3 images yielding the largest and smallest responses for each PC of interest.

## Discussion

Intermediate-level feature representations are an essential component of the human visual hierarchy, providing a link between low-level image properties and high-level conceptual information. We investigated this link by constructing voxelwise encoding models based on a set of image-computable texture statistics features (“P-S statistics”; Portilla and Simoncelli, 2000). In both early retinotopic and higher category-selective regions, our model generated accurate predictions of voxel responses to natural scene images that were held out during model fitting. Features contributing the most to the predictive accuracy of the model differed depending on position in the visual hierarchy: lower-level texture features explained more unique variance in early visual cortex, while higher-level texture features progressively explained a larger amount of unique variance in more anterior regions. At a finer-grained level, patterns of texture feature sensitivity were able to identify meaningful components of the overall representational space within visual cortex. These results increase our understanding of mid-level visual representations and highlight that such representations participate in processing even at later stages of the visual hierarchy.

Results broadly consistent with ours have been found in primate visual cortex. Multiple studies have shown that spectral features (i.e., the *energy-mean* features in our model), tend to have the largest regression model weights of any P-S feature type in V1, V2, and V4 (Hatanaka et al., 2022; Okazawa et al., 2017, 2015); this is consistent with our results showing the highest unique variance explained for the *energy-mean* features in every early visual ROI (Figure 8). Also consistent with Hatanaka et al. (2022), we found that the *pixel* features had the second-largest amount of unique variance explained in V1. More generally, our finding that higher-level feature sensitivity increased from V1 to hV4 – as well as increasing further in higher areas – is consistent with past studies reporting that the contributions of higher-order texture statistics are larger on average for V4 than either V1 (Hatanaka et al., 2022) or V2 (Okazawa et al., 2017). Importantly, our results expand upon these past findings by demonstrating the continuous increase in feature complexity beyond early visual cortex, and across a wide range of areas in the visual system.

Less aligned with past work is the relative importance of different higher-level texture feature subsets. In V1-hV4, we found that the higher-level texture feature subset which explained the most unique variance was the *linear-cross-orient* features. Although one study found relatively high average weights for this set of features (along with the *linear-cross-scale* features; Hatanaka et al., 2022), other studies did not include this feature subset in their models (Okazawa et al., 2017, 2015), and their results instead indicated a larger role for the *energy-cross-position* and *energy-cross-scale* features. These discrepancies may be the result of modeling differences across studies, such as the inclusion of certain feature subsets, particularly if some subsets carry redundant information with one another. Differences in stimulus type may also contribute to these discrepancies, as two studies (Okazawa et al., 2017, 2015) used synthetic texture stimuli as opposed to natural scene images, and sensitivity to higher-order texture features may be affected by the additional structure present in natural scene images. In particular, phase information may be important for modeling natural scene images, which are non-stationary across space, as compared to spatially homogeneous texture images. This is consistent with our observation that the *linear-cross-orient* and *linear-auto* features showed an advantage over the *energy-cross-orient* and *energy-auto* features (which are both phase-invariant). For the *linear-cross-orient* features, which contributed significant unique variance in early visual areas, higher-order selectivity may be driven by simple cells, whose responses are phase-dependent. Further work will be needed to evaluate these possibilities.

Although the lower-level features explained more unique variance in V1 than the higher-level features (Figure 5), a small but significant amount of unique variance in V1 in all 8 subjects was explained by the higher-level texture features (Table 5-1). In particular, the *linear-cross-orient* features, which include information about the relationships between different orientations in the image, explained a significant amount of variance in V1 (as well as V2, V3, and hV4). This result is somewhat surprising given that past work has found V1 neurons have little sensitivity to the higher-order correlation statistics of the P-S model (Freeman et al., 2013). However, other work suggests that V1 neurons are sensitive to higher-order statistical correlations (Purpura et al., 1994), and show context-dependent responses (e.g., figure-ground separation, sensitivity to illusory contours, or sensitivity for perceived size; Albright and Stoner, 2002; Carandini et al., 2005; Murray et al., 2006). Such contextual sensitivity may account for the observed selectivity to higher-order image structure in our results. Given the limited temporal resolution of fMRI, we cannot distinguish whether the sensitivity to higher-level texture statistics in V1 is due to feedforward processing, lateral interactions, or feedback from higher visual regions.

Sensitivity to higher-level texture features was also observed in more anterior areas, including voxels in and around category-selective visual regions such as EBA. When the contributions of fine-grained feature subsets were analyzed, the *linear-auto* texture features were among the most predictive subsets for higher visual areas, explaining a significant amount of unique variance at the ROI-averaged level in OPA, PPA, RSC, FFA, and EBA. This feature subset likely captures image periodicity, consistent with past work suggesting ventral visual areas VO1 and LOC are sensitive to the degree of spatial regularity in synthetic texture stimuli (Kohler et al., 2016). Other higher-level feature subsets also contributed significant amounts of unique variance in higher visual areas, with the *energy-cross-scale* features having the highest average unique variance in EBA. The cross-scale correlations included in these features may allow them to differentiate between different kinds of oriented elements, such as lines versus object edges, as well as capturing information about image scale (Figure 2, bottom left panel). The representation of these diverse sets of texture statistics features in higher visual cortex may support the broader computational functions of these regions.

Within higher visual cortex, higher-level texture model features explained relatively more unique variance in face- and body-selective areas (FFA, EBA), while lower-level model features contributed more in scene-selective areas (OPA, PPA, RSC). The sensitivity to lower-level features in scene-selective areas is consistent with work showing sensitivity to simple oriented features in scene-selective cortex (Lescroart et al., 2015; Nasr & Tootell, 2012), while the high sensitivity to higher-level features in face- and body-selective areas is consistent with the finding that the fusiform gyrus and the middle occipital gyrus exhibit larger responses to texture stimuli that include higher-order correlations (Beason-Held et al., 1998). This difference could also indicate that our model’s higher-level features align better with the mid-level features represented within face- and body-selective areas (curvy, organic features; Ponce et al., 2017) as compared to within scene selective areas (geometric, rectilinear features; Nasr et al., 2014). Processing in scene-selective cortical areas may also be more spatially global, reflecting a role in computing large-scale properties such as scene layout (Epstein & Baker, 2019). Our model features are computed over local regions of the image and, thus, may fail to capture the large-scale structure of scenes. Future work should improve the model in this direction.

Note that the overall performance of the model in more anterior regions was relatively poor compared to early areas, with *R*^2^ only reaching a fraction of the noise ceiling for most voxels. Thus, perhaps unsurprisingly, the texture statistics model does not fully capture the range of image features to which voxels in higher visual cortex are sensitive. This conclusion is consistent with past work suggesting that higher visual areas in macaques and humans, while exhibiting some sensitivity to P-S statistics, are also sensitive to certain high-level aspects of natural image structure that are not captured by these features (Long et al., 2018; Rust & DiCarlo, 2010). However, our results do indicate that intermediate visual features are reliably encoded within high-level visual areas, and that modeling these mid-level features can contribute to a more detailed understanding of the representations in these regions.

Relatedly, the ability of the texture statistics model to predict visual responses does not imply that the model features capture the explicit computations performed within these brain areas. As with any computational model, it is possible that the true features represented by the brain are merely correlated with the model features. Indeed, the similarity in performance between the texture statistics model and other low-level models suggests that there are multiple reasonable candidates. Nevertheless, the texture statistics model provides a simple and physiologically plausible hypothesis of how the brain computes mid-level feature representations, and we demonstrated that its features can predict voxel responses in visual cortex to natural images. Moreover, the contributions of different feature subsets to the model’s performance vary meaningfully across brain regions that serve different roles in visual processing. These findings pave the way for future work examining how mid-level features contribute to higher cognitive processes such as the recognition of complex objects and scenes.

## Acknowledgements

This research was funded by a Distinguished Postdoctoral Fellowship from the Carnegie Mellon Neuroscience Institute to MMH. Collection of the NSD dataset was supported by NSF IIS-1822683 and NSF IIS-1822929.

## Extended Data

Figure 3-1: View this analysis at: http://www.cs.cmu.edu/~mmhender/viewers/texturemodel/fullmodelR2/. This PyCortex webviewer shows the validation set accuracy of the texture statistics encoding model (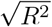) plotted on a cortical surface, for all 8 NSD subjects. White outlines and labels indicate the location of functionally-defined retinotopic and category-selective ROIs in each subject; see *Methods: Defining regions of interest (ROIs)* for details. Only voxels whose encoding model accuracy was above chance are shown (one-tailed *p*-values obtained using a permutation test, corrected for multiple comparisons, q=0.01.)

Figure 5-1: View this analysis at: http://www.cs.cmu.edu/~mmhender/viewers/texturemodel/varpart low vs high/. This PyCortex webviewer shows a comparison of the unique variance explained by the lower-level and higher-level texture features, for each of the 8 NSD subjects. The variance uniquely explained by lower-level and higher-level texture features was estimated using a variance partitioning analysis (see *Methods: Variance partitioning* for details). Values are in units of 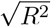.

Figure 7-1: View this analysis at: http://www.cs.cmu.edu/~mmhender/viewers/texturemodel/varpart all subsets/. This PyCortex webviewer shows the percentage of encoding model variance (units of 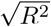) uniquely explained by each individual feature type, shown for each of the 8 NSD subjects. The maps in shades of blue correspond to subsets of the lower-level texture features, while maps in shades of red correspond to subsets of higher-level texture features. See *Methods: Variance partitioning* for details.

**Table 1-1:** Dimensionality of each subset of the texture statistics model features. First column shows the number of raw features computed in the model, other columns show the dimensionality of the higher-level model features following dimensionality reduction with PCA (see *Methods: Texture statistics features* for details); PCA was not used for the lower-level model features. Since the number of retained PCs could vary across pRFs, we report the median, minimum, and maximum number of PCs for each feature subset, across all pRFs.

|  | Raw | Median PCs | Min PCs | Max PCs |
| --- | --- | --- | --- | --- |
| pixel | 6 | - | - | - |
| energy-mean | 16 | - | - | - |
| linear-mean | 16 | - | - | - |
| marginal | 11 | - | - | - |
| energy-auto | 272 | 4 | 3 | 12 |
| linear-auto | 98 | 9 | 6 | 16 |
| energy-cross-orient | 24 | 5 | 4 | 6 |
| linear-cross-orient | 34 | 4 | 3 | 9 |
| energy-cross-scale | 48 | 7 | 5 | 12 |
| linear-cross-scale | 116 | 14 | 7 | 22 |

**Table 5-1:** First two rows show the number of individual subjects (out of 8 total) in which the unique variance explained by the lower-level and higher-level texture features was significantly greater than zero, for each ROI (one-tailed *p*-values computed using a bootstrap test; corrected for multiple comparisons; *q* =0.01; see *Methods: Variance partitioning* for details). Last row shows the number of subjects for which the difference between the unique variance explained by the lower- and higher-level texture features was significantly different (two-tailed *p*-values computed using a bootstrap test; corrected for multiple comparisons; *q* =0.01; see *Methods: Variance partitioning* for details). Where significant differences were detected, their signs were as follows: lower-level *>* higher-level in V1, V2, V3, hV4, PPA; higher-level *>* lower-level in EBA.

|  | V1 | V2 | V3 | hV4 | OPA | PPA | RSC | FFA | EBA |
| --- | --- | --- | --- | --- | --- | --- | --- | --- | --- |
| lower-level | 8 | 8 | 8 | 8 | 6 | 8 | 2 | 5 | 4 |
| higher-level | 8 | 8 | 7 | 5 | 8 | 8 | 3 | 7 | 8 |
| difference | 8 | 8 | 8 | 7 | 0 | 1 | 0 | 0 | 4 |

**Table 8-1:**
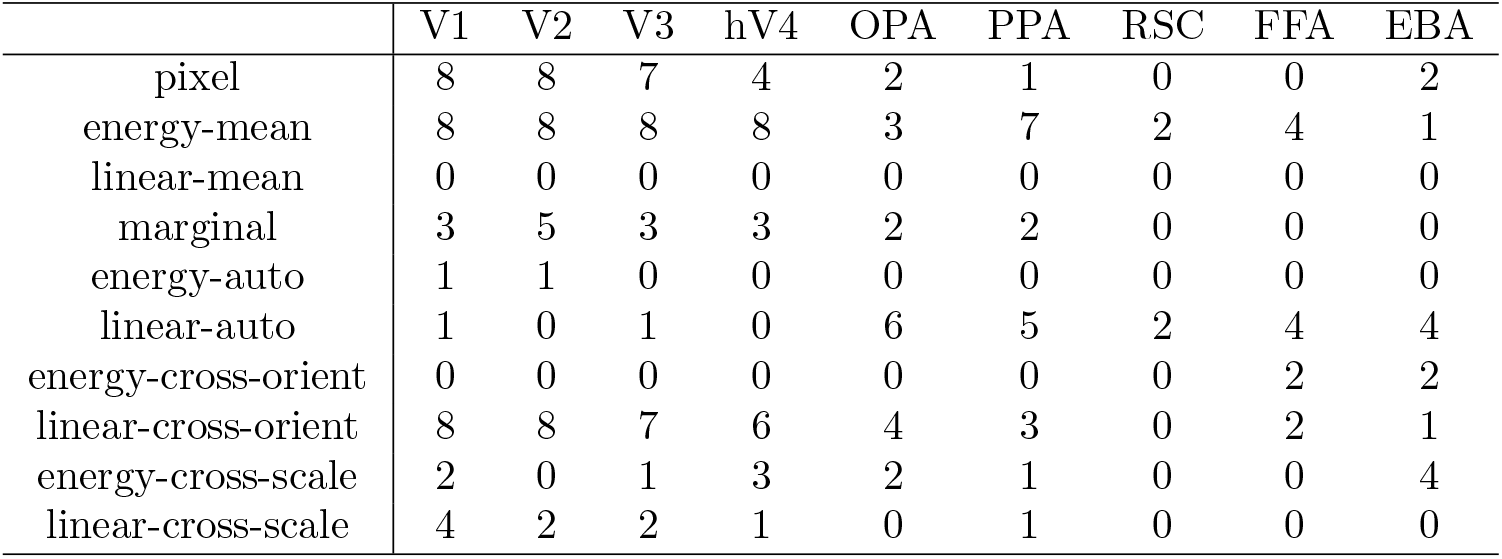
Number of individual subjects (out of 8 total) in which the unique variance explained by each subset of texture features was significantly greater than zero, for each ROI (one-tailed *p*-values computed using a bootstrap test; corrected for multiple comparisons; *q* =0.01; see *Methods: Variance partitioning* for details).

## Notes

### Competing Interest Statement

The authors have declared no competing interest.

### Summary of Updates

New analyses have been added.

## References

Albright, T. D., & Stoner, G. R. (2002). Contextual influences on visual processing. Annual Review of Neuroscience, 25, 339–379.

Allen, E. J., St-Yves, G., Wu, Y., Breedlove, J. L., Prince, J. S., Dowdle, L. T., Nau, M., Caron, B., Pestilli, F., Charest, I., Hutchinson, J. B., Naselaris, T., & Kay, K. (2021). A massive 7t fmri dataset to bridge cognitive neuroscience and artificial intelligence. Nature Neuroscience, 25, 116–126.

Balas, B. J. (2006). Texture synthesis and perception: Using computational models to study texture representations in the human visual system. Vision Research, 46, 299–309.

Balas, B. J., Nakano, L., & Rosenholtz, R. (2009). A summary-statistic representation in peripheral vision explains visual crowding. Journal of Vision, 9, 13–13.

Baumgartner, E., & Gegenfurtner, K. R. (2016). Image statistics and the representation of material properties in the visual cortex. Frontiers in Psychology, 7, 1185.

Beason-Held, L. L., Purpura, K. P., Krasuski, J. S., Maisog, J. M., Daly, E. M., Mangot, D. J., Desmond, R. E., Optican, L. M., Schapiro, M. B., & Vanmeter, J. W. (1998). Cortical regions involved in visual texture perception: A fmri study. Cognitive Brain Research, 7, 111–118.

Benjamini, Y., & Hochberg, Y. (1995). Controlling the false discovery rate: A practical and powerful approach to multiple testing. Journal of the Royal Statistical Society: Series B (Methodological), 57, 289–300.

Benson, N. C., Jamison, K. W., Arcaro, M. J., Vu, A. T., Glasser, M. F., Coalson, T. S., Essen, D. C. V., Yacoub, E., Ugurbil, K., Winawer, J., & Kay, K. (2018). The human connectome project 7 tesla retinotopy dataset: Description and population receptive field analysis. Journal of Vision, 18, 23–23.

Bergen, J. R., & Landy, M. S. (1991). Computational modeling of visual texture segregation. In M. S. Landy & J. A. Movshon (Eds.), Computational models of visual processing (pp. 253–271). The MIT Press.

Bracci, S., Ritchie, J., & de Beeck, H. O. (2017). On the partnership between neural representations of object categories and visual features in the ventral visual pathway. Neuropsychologia, 105, 153–164.

Carandini, M., Demb, J. B., Mante, V., Tolhurst, D. J., Dan, Y., Olshausen, B. A., Gallant, J. L., & Rust, N. C. (2005). Do we know what the early visual system does? The Journal of Neuroscience.

Connor, C. E., Brincat, S. L., & Pasupathy, A. (2007). Transformation of shape information in the ventral pathway. Current Opinion in Neurobiology, 17, 140–147.

Desimone, R., Albright, T. D., Gross, C. G., & Bruce, C. (1984). Stimulus-selective properties of inferior temporal neurons in the macaque. Journal of Neuroscience, 4, 2051–2062.

Dumoulin, S. O., & Wandell, B. A. (2008). Population receptive field estimates in human visual cortex. NeuroImage, 39, 647–660.

Duncan, R. O., & Boynton, G. M. (2003). Cortical magnification within human primary visual cortex correlates with acuity thresholds. Neuron, 38, 659–671.

Epstein, R. A., & Baker, C. I. (2019). Scene perception in the human brain. Annual review of vision science, 5, 373–397.

Freeman, J., Brouwer, G. J., Heeger, D. J., & Merriam, E. P. (2011). Orientation decoding depends on maps, not columns. Journal of Neuroscience, 31, 4792–4804.

Freeman, J., & Simoncelli, E. P. (2011). Metamers of the ventral stream. Nature Neuroscience, 14, 1195–1204.

Freeman, J., Ziemba, C. M., Heeger, D. J., Simoncelli, E. P., & Movshon, J. A. (2013). A functional and perceptual signature of the second visual area in primates. Nature Neuroscience, 16, 974–981.

Gao, J. S., Huth, A. G., Lescroart, M. D., & Gallant, J. L. (2015). Pycortex: An interactive surface visualizer for fmri. Frontiers in Neuroinformatics, 9.

Grill-Spector, K., & Weiner, K. S. (2014). The functional architecture of the ventral temporal cortex and its role in categorization. Nature Reviews Neuroscience, 15, 536–548.

Groen, I. I. A., Silson, E. H., & Baker, C. I. (2017). Contributions of low- and high-level properties to neural processing of visual scenes in the human brain. Philosophical Transactions of the Royal Society B: Biological Sciences, 372.

Groen, I. I., Ghebreab, S., Lamme, V. A., & Scholte, H. S. (2012). Spatially pooled contrast responses predict neural and perceptual similarity of naturalistic image categories. PLOS Computational Biology, 8, e1002726.

Güçlü, U., & van Gerven, M. A. (2014). Unsupervised feature learning improves prediction of human brain activity in response to natural images. PLOS Computational Biology, 10, e1003724.

Hatanaka, G., Inagaki, M., Takeuchi, R. F., Nishimoto, S., Ikezoe, K., & Fujita, I. (2022). Processing of visual statistics of naturalistic videos in macaque visual areas v1 and v4. Brain Structure and Function, 227, 1385–1403.

Henderson, M., Tarr, M. J., & Wehbe, L. (2022). Low-level tuning biases in higher visual cortex reflect the semantic informativeness of visual features. bioRxiv.

Holm, S. (1979). A simple sequentially rejective multiple test procedure. Scandinavian journal of statistics, 65–70.

Hubel, D. H., & Wiesel, T. N. (1962). Receptive fields, binocular interaction and functional architecture in the cat’s visual cortex. The Journal of Physiology, 160, 106.

Huth, A. G., Heer, W. A. D., Griffiths, T. L., Theunissen, F. E., & Gallant, J. L. (2016). Natural speech reveals the semantic maps that tile human cerebral cortex. Nature, 532, 453–458.

Kohler, P. J., Clarke, A., Yakovleva, A., Liu, Y., & Norcia, A. M. (2016). Representation of maximally regular textures in human visual cortex. Journal of Neuroscience, 36, 714–729.

Konkle, T., & Caramazza, A. (2013). Tripartite organization of the ventral stream by animacy and object size. Journal of Neuroscience, 33, 10235–10242.

Krizhevsky, A. (2014). One weird trick for parallelizing convolutional neural networks.

Lescroart, M. D., & Gallant, J. L. (2019). Human scene-selective areas represent 3d configurations of surfaces. Neuron, 101, 178–192.e7.

Lescroart, M. D., Stansbury, D. E., & Gallant, J. L. (2015). Fourier power, subjective distance, and object categories all provide plausible models of bold responses in scene-selective visual areas. Frontiers in Computational Neuroscience, 9, 1–20.

Lin, T. Y., Maire, M., Belongie, S., Hays, J., Perona, P., Ramanan, D., Dollár, P., & Zitnick, C. L. (2014). Microsoft coco: Common objects in context. Lecture Notes in Computer Science (including subseries Lecture Notes in Artificial Intelligence and Lecture Notes in Bioinformatics), 8693 LNCS, 740–755.

Long, B., Konkle, T., Cohen, M. A., & Alvarez, G. A. (2016). Mid-level perceptual features distinguish objects of different real-world sizes. Journal of Experimental Psychology: General, 145, 95–109.

Long, B., Störmer, V. S., & Alvarez, G. A. (2017). Mid-level perceptual features contain early cues to animacy. Journal of Vision, 17, 20.

Long, B., Yu, C. P., & Konkle, T. (2018). Mid-level visual features underlie the high-level categorical organization of the ventral stream. Proceedings of the National Academy of Sciences of the United States of America, 115, E9015–E9024.

Murray, S. O., Boyaci, H., & Kersten, D. (2006). The representation of perceived angular size in human primary visual cortex. Nat Neurosci, 9 (3), 429–434.

Naselaris, T., & Kay, K. N. (2015). Resolving ambiguities of mvpa using explicit models of representation. Trends in Cognitive Sciences, 19, 551–554.

Nasr, S., Echavarria, C. E., & Tootell, R. B. (2014). Thinking outside the box: Rectilinear shapes selectively activate scene-selective cortex. Journal of Neuroscience, 34, 6721–6735.

Nasr, S., & Tootell, R. B. (2012). A cardinal orientation bias in scene-selective visual cortex. Journal of Neuroscience, 32, 14921–14926.

Nunez-Elizalde, A. O., Huth, A. G., & Gallant, J. L. (2019). Voxelwise encoding models with non-spherical multivariate normal priors. NeuroImage, 197, 482–492.

Okazawa, G., Tajima, S., & Komatsu, H. (2017). Gradual development of visual texture-selective properties between macaque areas v2 and v4. Cerebral Cortex, 27, 4867–4880.

Okazawa, G., Tajima, S., & Komatsu, H. (2015). Image statistics underlying natural texture selectivity of neurons in macaque v4. Proceedings of the National Academy of Sciences of the United States of America, 112, E351–E360.

Oliva, A., & Torralba, A. (2001). Modeling the shape of the scene: A holistic representation of the spatial envelope *. International Journal of Computer Vision, 42, 145–175.

Peirce, J. W. (2015). Understanding mid-level representations in visual processing. Journal of Vision, 15, 5–5.

Ponce, C. R., Hartmann, T. S., & Livingstone, M. S. (2017). End-stopping predicts curvature tuning along the ventral stream. Journal of Neuroscience, 37, 648–659.

Portilla, J., & Simoncelli, E. P. (2000). Parametric texture model based on joint statistics of complex wavelet coefficients. International Journal of Computer Vision, 40, 49–71.

Prince, J. S., Charest, I., Kurzawski, J. W., Pyles, J. A., Tarr, M. J., & Kay, K. N. (2022). Glmsingle: A toolbox for improving single-trial fmri response estimates. bioRxiv, 2022.01.31.478431.

Purpura, K. P., Victor, J. D., & Katz, E. (1994). Striate cortex extracts higher-order spatial correlations from visual textures. Proceedings of the National Academy of Sciences of the United States of America, 91, 8482–8486.

Rosenholtz, R., Huang, J., Raj, A., Balas, B. J., & Ilie, L. (2012). A summary statistic representation in peripheral vision explains visual search. Journal of Vision, 12, 14–14.

Roth, Z. N., Kay, K., & Merriam, E. P. (2022). Natural scene sampling reveals reliable coarse-scale orientation tuning in human v1. Nature Communications 2022 13:1, 13, 1–13.

Rust, N. C., & DiCarlo, J. J. (2010). Selectivity and tolerance (“invariance”) both increase as visual information propagates from cortical area v4 to it. Journal of Neuroscience, 30, 12978–12995.

Serences, J. T., & Saproo, S. (2012). Computational advances towards linking bold and behavior. Neuropsychologia, 50, 435–446.

Simoncelli, E. P., & Freeman, W. T. (1995). Steerable pyramid: A flexible architecture for multi-scale derivative computation. IEEE International Conference on Image Processing, 3, 444–447.

Stigliani, A., Weiner, K. S., & Grill-Spector, K. (2015). Temporal processing capacity in high-level visual cortex is domain specific. Journal of Neuroscience, 35, 12412–12424.

St-Yves, G., & Naselaris, T. (2017). The feature-weighted receptive field: An interpretable encoding model for complex feature spaces. NeuroImage, 180, 188–202.

Ullman, S., Vidal-Naquet, M., & Sali, E. (2002). Visual features of intermediate complexity and their use in classification. Nature Neuroscience, 5, 682–687.

Walther, D. B., & Shen, D. (2014). Non-accidental properties underlie human categorization of complex natural scenes. Psychological science, 25, 851.

Wang, L., Mruczek, R. E., Arcaro, M. J., & Kastner, S. (2015). Probabilistic maps of visual topography in human cortex. Cerebral Cortex, 25, 3911–3931.

Wehbe, L., Murphy, B., Talukdar, P., Fyshe, A., Ramdas, A., & Mitchell, T. (2014). Simultaneously uncovering the patterns of brain regions involved in different story reading subprocesses. PLOS ONE, 9, e112575.

Wu, M. C., David, S. V., & Gallant, J. L. (2006). Complete functional characterization of sensory neurons by system identification. https://doi.org/10.1146/annurev.neuro.29.051605.113024, 29, 477–505.

